# Low heritability and high phenotypic plasticity of salivary cortisol in response to environmental heterogeneity in a wild pinniped

**DOI:** 10.1101/2021.10.08.463664

**Authors:** Rebecca Nagel, Sylvia Kaiser, Claire Stainfield, Camille Toscani, Cameron Fox-Clarke, Anneke J. Paijmans, Camila Costa Castro, David L. J. Vendrami, Jaume Forcada, Joseph I. Hoffman

**Author notes:** Joint senior authors (Jaume Forcada, Joseph I. Hoffman). Corresponding author: Rebecca Nagel, Department of Animal Behaviour, Bielefeld University, 33501 Bielefeld, Germany.

## Abstract

Individuals are unique in how they interact with and respond to their environment. Correspondingly, unpredictable challenges or environmental stressors often produce an individualized response of the hypothalamic-pituitary-adrenal axis and its downstream effector cortisol. We used a fully crossed, repeated measures design to investigate the factors shaping individual variation in baseline cortisol in Antarctic fur seal pups and their mothers. Saliva samples were collected from focal individuals at two breeding colonies, one with low and the other with high density, during two consecutive years of contrasting food availability. Mothers and pups were sampled concurrently at birth and shortly before weaning, while pups were additionally sampled every 20 days. We found that heritability was low for baseline cortisol, while within-individual repeatability and among-individual variability were high. A substantial proportion of the variation in baseline cortisol could be explained in pups and mothers by a combination of intrinsic and extrinsic factors including sex, weight, day, season, and colony of birth. Our findings provide detailed insights into the individualization of endocrine phenotypes and their genetic and environmental drivers in a wild pinniped. Furthermore, the strong associations between cortisol and life history traits that we report in fur seals could have important implications for understanding the population dynamics of species impacted by environmental change.

## Introduction

Despite similarities in age, sex, or social status, individuals often differ in how they interact with their environment (Réale and Dingemanse 2010; Dall et al. 2012). Once perceived as a statistical nuisance, an ever growing body of evidence now suggests that these individual differences are often consistent and stable across time and contexts, with profound implications for understanding phenotypic variation, niche specialization, and animal personality (Sih et al. 2012; Wolf and Weissing 2012). The past decade has thus witnessed an increased awareness of individualization as a fundamentally important and compelling aspect of evolutionary biology, ecology, and animal behavior (Bolnick et al. 2003; Trillmich et al. 2018; Krüger et al. 2021).

Consistent differences among individuals are likely to be mediated by a combination of intrinsic and extrinsic factors and can be understood as the interaction between an individual’s phenotype, genotype, and its ecological context such that its fitness is maximized (Dingemanse and Réale 2005; Vessey and Drickamer 2010; Lihoreau et al. 2021). At the proximate level, individualized phenotypic adjustments to environmental factors may be governed by, among other things, variation in the concentrations of circulating hormones (Müller et al. 2020). In particular, individual variation in cortisol, an important hormone in the physiological stress response, appears to play a major role in shaping individual responses to environmental conditions (Wingfield and Romero 2011).

Cortisol is a steroid hormone that belongs to the class of glucocorticoids. Its release is regulated by the hypothalamic-pituitary-adrenal (HPA) axis. Glucocorticoids play an essential role in maintaining metabolic and homeostatic functions (Kuo et al. 2015). Under predictable conditions, cortisol is released continuously at baseline levels that vary naturally throughout the day and over an individual’s lifetime (Lightman and Conway-Campbell 2010). In the face of unpredictable challenges, however, activation of the HPA axis results in increased levels of secreted cortisol for the duration of the stressor, before levels return to baseline (Bellavance and Rivest 2014).

While the physiology of the HPA axis is largely conserved across mammals (Romero and Butler 2007; Wingfield and Romero 2011), empirical studies have documented significant differences in cortisol concentrations among species and individuals (Schoenemann and Bonier 2018; Taff et al. 2018), as well as notable variation in the HPA axis response across time and space (Romero and Gormally 2019). Such phenotypic variation in response to intrinsic and environmental differences could potentially facilitate adaptation to different habitats and conditions (Sih et al. 2012). For example, cortisol levels have been shown to vary significantly among individuals experiencing different densities (Meise et al. 2016) and levels of nutritional stress (Kitaysky et al. 2007). Variation in cortisol among individuals has also been linked to intrinsic factors such as age (Pavitt et al. 2015), sex (Azevedo et al. 2019), and weight (Jeanniard du Dot et al. 2009). Nonetheless, when such factors are accounted for, empirical studies have shown that cortisol levels are often highly repeatable within individuals (Schoenemann and Bonier 2018; Taff et al. 2018). In guinea pigs, for example, cortisol responsiveness and to a lesser extent baseline cortisol levels are highly repeatable from late adolescence to adulthood (Mutwill et al. 2021).

Given that cortisol levels often exhibit high among-individual variability and within-individual repeatability, it has been argued that genetic rather than environmental factors could explain much of the observed phenotypic variance, which would imply a high evolvability of this endocrine trait (Boake 1989; Jenkins et al. 2014). Correspondingly, several empirical studies have reported moderate to high levels of cortisol heritability in free-living vertebrate populations (Jenkins et al. 2014; Stedman et al. 2017; Bairos-Novak et al. 2018). However, persistent intrinsic and / or extrinsic factors might also produce repeatable phenotypes regardless of the underlying genotype (Taff et al. 2018), a possibility that has often been overlooked in the literature (Bonier and Martin 2016). This is particularly true when genetically similar individuals experience a similar temporal and spatial environment leading to, for example, phenotypic similarity in the endocrine phenotype in response to current internal and environmental stimuli (contextual plasticity) or stimuli encountered in the past (developmental plasticity) (Stamps and Biro 2016). If unaccounted for, such phenotype-environment correlations may upwardly bias the estimated additive genetic variance for phenotypic traits (Kruuk et al. 2003).

Pinnipeds, and otariids in particular, are ideally suited to investigate the effects of internal and environmental factors on cortisol levels. First, otariids are colonially breeding, with males competing to establish and maintain harems on densely packed breeding beaches (Forcada and Staniland 2018). Cortisol may play an important role in how individuals adapt to this dynamic environment by restoring homeostasis after unpredictable challenges such as territorial bouts or unwanted mating attempts. Second, while many breeding beaches do not differ appreciably in qualities such as substrate type or topology, the density of individuals often varies from one place to another, setting up a spatial dynamic that tends to remain stable over time (Cassini 1999). Consequently, as pups are born on land and remain ashore throughout much of their early ontogeny (Payne 1979; McCafferty et al. 1998), cortisol might play an important role in mediating individual responses to variation in density. Finally, cortisol levels have been investigated in several pinniped species in relation to ontogeny (Ortiz et al. 2003; Atkinson et al. 2011), environmental conditions (DeRango et al. 2019), and handling regimes (Engelhard et al. 2002; Harcourt et al. 2010; Bennett et al. 2012; Champagne et al. 2012). Methodologies for collecting and assessing cortisol in pinnipeds are therefore well established in the literature.

Our model otariid species, the Antarctic fur seal (*Arctocephalus gazella*), has been extensively studied by the British Antarctic Survey (BAS) on Bird Island, South Georgia since the 1980s. Two breeding colonies on the island provide a unique “natural experiment” for investigating individual responses to population density. Freshwater Beach (FWB) and Special Study Beach (SSB) are situated less than 200 meters apart (Figure 1a), meaning they are exposed to comparable climatic conditions. Breeding females from both locations also likely forage in the same areas (Hunt et al. 1992) and do not differ significantly in quality traits such as body size and condition (Nagel et al. 2021). Despite these similarities, the two colonies differ in the density of conspecifics. Direct counts of individuals ashore suggest that the density of breeding females is almost four times higher at SSB than FWB (Meise et al. 2016) and the modal local density of focal pups across the entire breeding season is also higher for pups born at SSB (Nagel et al. 2021).

**Figure 1:**
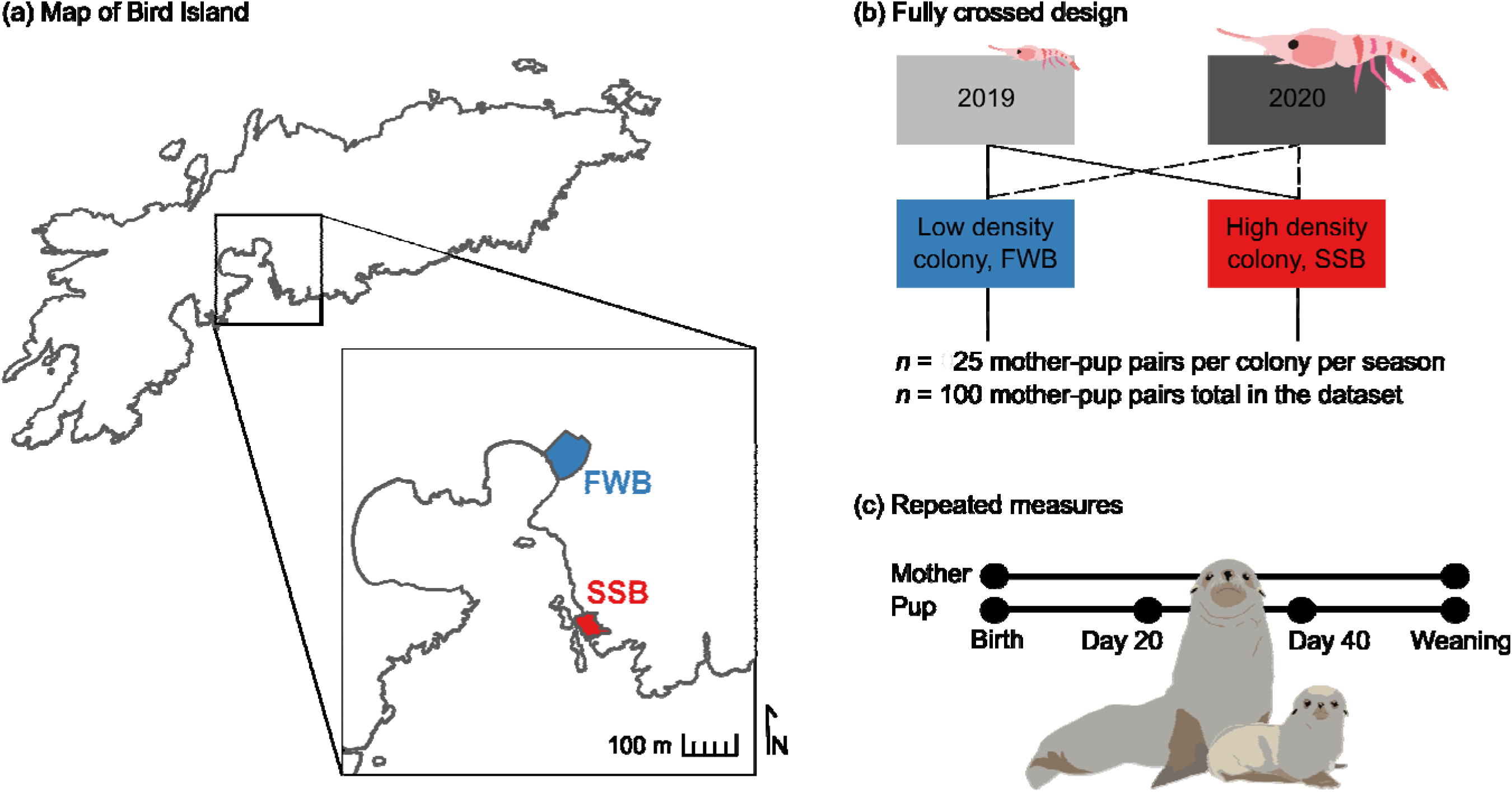
Location and study design. (a) Map of Bird Island, South Georgia, a sub-Antarctic island in the southern Atlantic Ocean. The inset shows an enlarged view of the two study colonies from which mother-pup pairs were sampled. Freshwater Beach (FWB, shown in blue) and Special Study Beach (SSB, shown in red) are separated by approximately 200 meters. (b) We employed a fully crossed sampling scheme involving the collection of saliva samples from a total of 100 pairs from the two colonies in two successive breeding seasons, the first of which was coincidentally a year of particularly low food availability. (c) Each focal mother was sampled twice in a season while pups were sampled every 20 days from birth until weaning.

We took advantage of this unique natural setup to investigate the intrinsic and extrinsic factors shaping individual variation in baseline cortisol in Antarctic fur seal pups and their mothers. We used a fully crossed, repeated measures design (Figure 1b, c) comprising longitudinal data from mother-offspring pairs from the two colonies over two consecutive breeding seasons, the first of which was coincidentally one of the worst years on record with respect to food availability (Nagel et al. 2021). Specifically, we collected saliva samples and accompanying biometric data from 25 randomly selected focal pairs from each colony in both seasons. To quantify baseline cortisol levels, saliva was collected immediately, within three minutes of capture. Mothers were sampled twice during the breeding season, while pups were sampled every 20 days from birth until just before molting at around 60 days of age.

We used animal models to obtain heritability estimates for baseline cortisol using both a simple pedigree and a genomic relatedness matrix obtained from a high density single nucleotide polymorphism (SNP) array (Humble et al. 2020). We then used linear mixed models to evaluate the within- and among-individual variability of cortisol levels in pups and their mothers. Included in each model as explanatory variables were multiple intrinsic (sex, weight, body condition, and days after initial sampling) and extrinsic (density and year) variables. In line with previous studies of wild vertebrate populations, we hypothesized that cortisol levels would be heritable and repeatable within individuals. We further hypothesized that baseline cortisol would be higher in pups and mothers from the high-density colony and in the season of low food availability.

## Materials and Methods

### Field study

This study was conducted during the Antarctic fur seal breeding seasons (December to March) of 2018–19 (hereafter 2019) and 2019–20 (hereafter 2020) at Bird Island, South Georgia (54°00’24.8□ S, 38°03’04.1□ W). Each season, we sampled 25 unique mother-pup pairs from two neighboring breeding colonies, one of low (FWB) and the other of high (SSB) density (Figure 1a). Sampling at both locations was randomized with respect to pup sex, resulting in a final sample size of 51 male and 49 female pups. Pup mortality was higher at FWB than SSB (32% *versus* 12%, respectively), and averaged 25.6% over the two colonies and seasons.

Each mother and her pup were captured concurrently on two separate occasions: 2–3 days postpartum (December) and again as the pups began to moult shortly before weaning (March). Pups were additionally recaptured every 20 days. For the capture, restraint, and sampling of individuals, we employed protocols that have been established and refined over 30 consecutive years of the BAS long-term monitoring and survey program. Briefly, adult females were captured with a noosing pole and held on a restraint board during processing. Pups were captured with a slip noose or by hand and were restrained by hand. After sampling, individuals were released as close to their capture site as possible and, when present, pups were reunited with their mothers.

At first sampling, focal individuals were fitted with VHF transmitters to the dorsal side of the neck between the shoulder blades with epoxy glue (pups: Sirtrack core marine glue-on V2G 152A; mothers: Sirtrack core marine glue-on V2G 154C). Transmitter signals were monitored throughout the season using a hand-held VHF receiver (AOR LTD., AR8200). Focal individuals were also given cattle ear tags (Dalton Supplies, Henley on Thames, UK) in the trailing edge of each foreflipper (Gentry and Holt 1982) for identification. Tissue plugs were collected and stored in 20% dimethyl sulfoxide (DMSO) saturated with salt at −20°C for subsequent genetic analysis.

At every capture, weight and length measurements were taken from which a scaled mass index was calculated according to (Peig and Green 2009). This condition metric serves as a reliable indicator of overall fitness as it has been correlated with, among other things, offspring survival (Milenkaya et al. 2015; Gélin et al. 2016) and mating success (Gastón and Vaira 2020). At every capture, the saliva sample was taken within three minutes of capture to provide data on baseline cortisol levels (Bozovic et al. 2013). Saliva was collected by rotating sterile cotton tip applicators fitted in polypropylene tubes (ROTH, Art. No. XC10.2) in the cheek pouch and under the tongue. Samples were centrifuged and stored at −20°C for subsequent cortisol analysis.

### Hormone quantification

Saliva samples were thawed and centrifuged for ten minutes to separate the mucins. The clear supernatant was then used for the determination of cortisol concentrations. Samples contaminated with blood (reddish supernatant) were discarded (*n* = 30 in 2019 and *n* = 31 in 2020), as cortisol values are often falsely elevated in such samples. Cortisol concentrations were determined in duplicate using enzyme-linked immunosorbent assays (cortisol free in saliva DES6611, Demeditec Diagnostics GmbH, Kiel, Germany). We calculated the average coefficient of variation (CV) resulting from individual CVs for all duplicates in an assay. Mean intra-assay CV for a total of 12 assays was 3.89%. All samples were determined in duplicate and if CV was larger than 10%, determination of the sample was repeated. Furthermore, two samples of different concentrations were run in duplicate on a total of 12 plates to assess inter-assay variation, which was on average 4.36%. The antibody showed the following cross-reactivities: cortisol 100%, 11-desoxycortisol 50%, corticosterone 6.2%, 11-desoxycorticosterone 2.6%, 17α-oh-progesterone 1.3%, cortisone and prednisone < 1%, testosterone, estradiol, and androstendione < 0.1%. For additional information on kit validation as per the linearity and recovery rate, see the respective Supplementary Tables S1 and S2.

### SNP genotyping and genomic relatedness matrix construction

For the 95 focal individuals sampled in 2019, we extracted total genomic DNA from tissue samples using a standard chloroform-isoamylalcohol protocol (for a description of the full protocol, see the Supplementary Materials). SNP genotyping was performed on these samples using a custom Affymetrix SNP array as described by (Humble et al. 2020). Quality control of the raw output data and genotyping were implemented using the Axiom Analysis Suite (5.0.1.38, Affymetrix) based on parameter thresholds set to their default values for diploid organisms. SNPs initially classified as “off target variants” (OTV) were recovered using the “Run OTV caller” function. Of the 85,359 SNPs tiled on the array, 77,873 were retained for further analysis representing SNPs classified as “PolyHighResolution” (SNPs that passed all of the Axiom Analysis Suite quality controls) and “NoMinorHomozygote” (SNPs that passed all quality controls but no homozygote genotypes for the minor allele were found). An additional 3,423 SNPs with minor allele frequencies below 0.01 and 2,096 SNPs that departed significantly from Hardy Weinberg equilibrium (HWE) were removed using PLINK version 1.9 (Purcell et al. 2007). Departures from HWE were identified based on an alpha level of 0.01 after implementing mid-*p* adjustment (Graffelman and Moreno 2013). After filtering, a total of 72,354 SNPs were retained and used to produce a genomic relatedness matrix using the --make-grm option in GCTA version 1.93.1 (Yang et al. 2011).

### Heritability of cortisol levels

To quantify the proportion of the total variance in baseline cortisol attributable to genetic differences among individuals, we fitted two multivariate generalized linear mixed models (GLMMs) in *MCMCglmm* (Hadfield 2010) with baseline cortisol as the dependent variable and individual ID and relatedness as random effects. For the first model, a simple pedigree (comprising mother-offspring pairs) was built for the full dataset providing an estimate of maternal genetic effects on cortisol concentrations. The second model incorporated the SNP relatedness matrix from individuals sampled only in the first season (2019) and, although smaller in sample size, provides a less biased estimate of genetic relatedness by including not only mother-offspring pairs but also detecting, for example, half-siblings. We used weak but informative priors (0.05 of the observed phenotypic variance) in both models. Markov chains were run for 9,000,000 iterations and we retained every 8,500^th^ value after removing 150,000 iterations of burn-in to generate posterior distributions of the random parameters. The posterior distribution of the model intercept and autocorrelation were checked to assess model fit. We obtained estimates of baseline cortisol heritability by dividing the additive genetic variance by the total phenotypic variance (*h*^2^ = V_A_/V_P_) for each sample of the posterior distribution.

### Estimating intrinsic and extrinsic factors influencing baseline cortisol

To determine whether an individual’s weight, body condition, age, sex, season, colony, and an interaction between season and colony explained a significant proportion of the variation in baseline cortisol among pups, we fitted a GLMM with a log-link gamma error distribution in *lme4* (Bates et al. 2015). A second GLMM with maternal cortisol as the dependent variable was used to determine the proportion of variation explained by an individual’s weight, body condition, the number of days postpartum, season, colony, and an interaction between season and colony. To account for both structural and data multicollinearity among weight, condition, and age (pups) / days postpartum (mothers), these variables were rescaled and centered by subtracting the mean from all observed values. In preliminary analyses, we tested for the presence of heterogeneous variance by allowing individual slopes to vary by age (pups) or days postpartum (mothers). For both models, random intercepts were included for each individual to account for repeated measures. The residuals of the full models were inspected for linearity and equality of error variances (using plots of residuals versus fits), normality (using Q – Q plots), and homogeneity of variance (using Levene’s test). A backward elimination based on the chi-squared statistic was implemented to simplify the full models such that, in each iteration, the fixed effect with the lowest chi-squared value was removed from the model until we only tested the null model. The best fitting models were then taken to be those with the lowest AIC values. We present only the best model from each analysis in the Results, but model reduction and AIC scores for all models are available in the Supporting R Markdown file. The statistical significance of fixed predictors was assessed using Wald tests. We determined the marginal *R*^2^ (variance explained by fixed effects) and conditional *R*^2^ (variance explained by fixed and random effects) according to (Nakagawa and Schielzeth 2013).

### Animal handling, ethics and permits

Sampling was carried out by the BAS under permits from the Government of South Georgia and the South Sandwich Islands (Wildlife and Protected Areas Ordinance (2011), RAP permit numbers 2018/024 and 2019/032). The samples were imported into the UK under permits from the Department for Environment, Food, and Rural Affairs (Animal Health Act, import license number ITIMP18.1397) and from the Convention on International Trade in Endangered Species of Wild Fauna and Flora (import numbers 578938/01-15 and 590196/01-18). All procedures used were approved by the BAS Animal Welfare and Ethics Review Body (AWERB applications 2018/1050 and 2019/1058).

## Results

We used a fully crossed, repeated measures design incorporating saliva samples from 96 unique pups and 93 unique mothers from two colonies of contrasting density across two consecutive years of contrasting food availability (Figure 1). Sample sizes were balanced between the colonies (*n* = 93 from FWB and *n* = 96 from SSB) and seasons (*n* = 95 from 2019 and *n* = 94 from 2020). Each season, pups were sampled every 20 days from birth until weaning, amounting to a total of 290 analyzed saliva samples. Mothers were sampled twice each season, once shortly after birth and again shortly before molting, which amounted to a total of 145 analyzed saliva samples.

### Cortisol heritability estimates

We estimated heritability of baseline cortisol using two animal models, the first incorporating known pedigree relationships (i.e. mother-offspring pairs from both years) and the second incorporating a SNP relatedness matrix, which was only available for the first year of the study. Narrow-sense heritability (*h*^2^) estimates from both models were low with overlapping 95% credible intervals (pedigree model: *h*^2^ = 0.013, 95% highest posterior density 0.004 – 0.045; SNP relatedness model: *h^2^* = 0.018, 95% highest posterior density 0.004 – 0.062) (Figure 2). The additive genetic (V_A_) and residual (V_R_) variance estimates of the two models were also comparable (pedigree model: 95% highest posterior density of V_A_ = 0.1 – 1.6 and V_R_ = 30.6 – 39.9; SNP relatedness model: 95% highest posterior density of V_A_ = 0.1 – 2.3 and V_R_ = 28.5 – 41.7).

**Figure 2:**
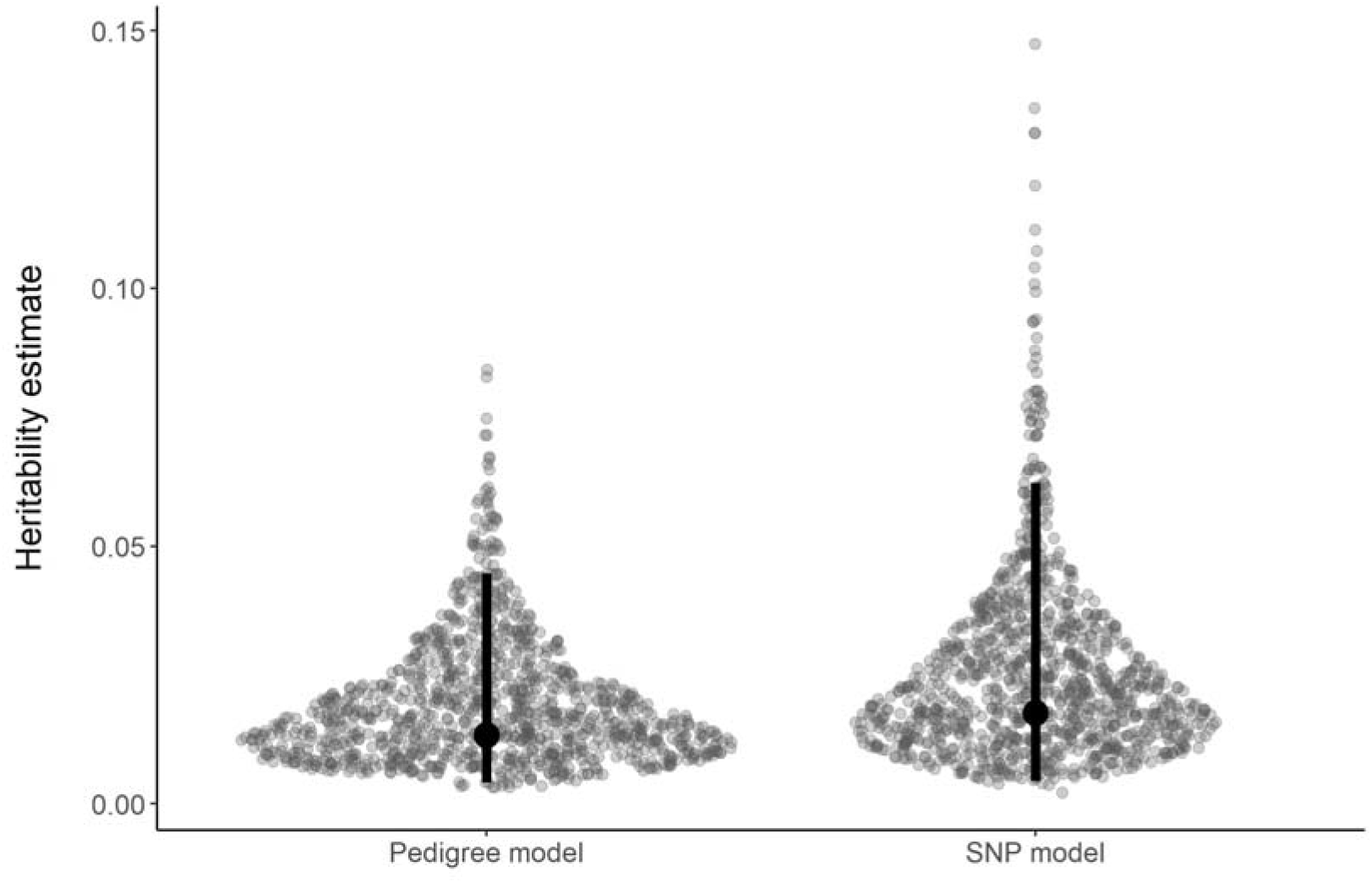
Posterior distributions of heritability (*h*^2^) estimates for baseline cortisol. A simple pedigree (mother-offspring pairs) for the entire dataset and a custom 85K SNP array for the 95 individuals sampled in the 2019 season were used to calculate the relatedness matrix. The modes and highest posterior density intervals of the posterior distributions are shown as points and bars, respectively.

### Intrinsic and extrinsic factors influencing baseline cortisol

The best supported model of pup baseline cortisol contained individual age (*p* < 0.001), weight (*p* < 0.001), sex (*p* = 0.004), season (*p* = 0.003), body condition (*p* = 0.093), and colony (*p* = 0.107) as fixed effects (Table 1a, Figure 3a). The total amount of variance explained by this model was high (conditional *R*^2^ = 0.657), as was the repeatability of baseline cortisol values across individuals (ICC = 0.39). Including ID as a random effect significantly improved the fit of the model, indicating appreciable among-individual variability in baseline cortisol (*p* < 0.001; Supplementary Table S1a). Allowing individual slopes to vary between age groups also significantly improved model fit (*p* < 0.001; Supplementary Table S3) suggesting that individuals responded to the covariates differently depending on their age. Overall, baseline cortisol decreased with increasing pup age (Figure 3b) and was higher among pups born in 2020, the year of higher food availability (Figure 3c). Baseline cortisol decreased significantly as pup weight increased (Figure 3d), although the slope of the regression between cortisol and weight approached zero as pups approached their moult. Finally, baseline cortisol tended to be higher in males than females (Figure 3e).

**Figure 3:**
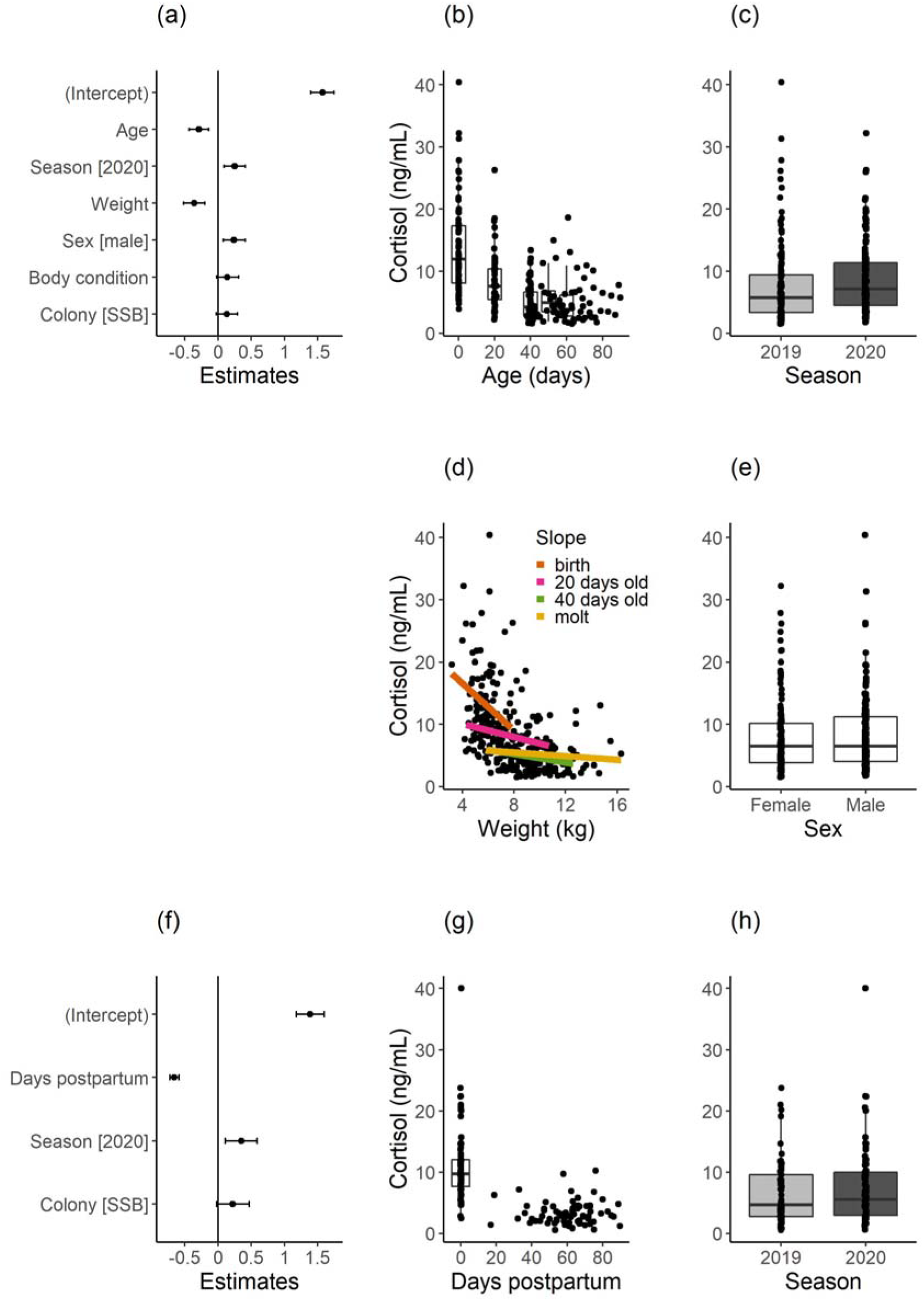
Generalized linear mixed models for pup (a – e) and mother (f – h) baseline cortisol values. Estimates ± 95% confidence intervals for all fixed effects included in the best fit models for pups and mothers are shown in panels a and f, respectively. Significant main effects for both models are shown in panels b – e and panels g – h, respectively. Boxes show median values ± 75% percentiles with the vertical lines indicating 95% confidence intervals. Further details of the model output can be found in Table 1.

**Table 1:**
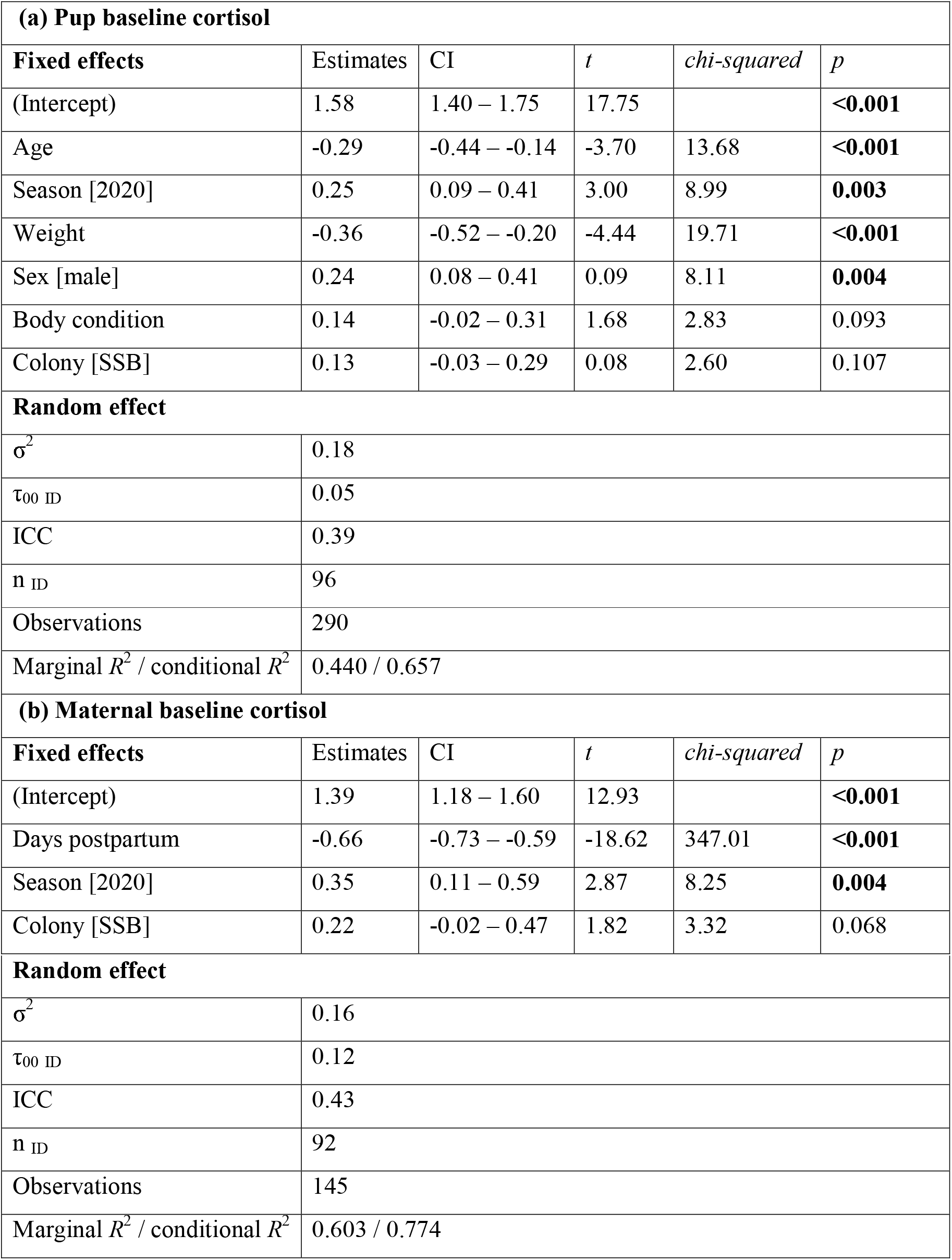
Parameter estimates from the best fit generalized linear mixed models of (a) pup and (b) maternal baseline cortisol. Random intercepts were included for each individual to account for repeated measures. Estimates together with their 95% confidence intervals (CI) as well as Wald *t*-values and chi-squared values are presented. Significant *p*-values are in bold. The mean squared error (σ^2^), between group variance (τ00), Intraclass Correlation Coefficient (ICC; the consistency within an individual across multiple measurements), the sample size (*n*) and total number of observations, as well as the variance explained by the fixed effects (marginal *R*^2^) and variance explained by both fixed and random effects (conditional *R*^2^) are given.

The best-supported model of maternal baseline cortisol contained days postpartum (*p* < 0.001), season (*p* = 0.004), and colony (*p* = 0.068) as fixed effects (Table 1b, Figure 3f). Neither individual weight nor body condition were retained in the model. The total variance explained by the model was again high (conditional *R*^2^ = 0.774), as was repeatability of baseline cortisol within individuals (ICC = 0.43). Including ID as a random effect significantly improved model fit, indicating appreciable among-individual variability in baseline cortisol (*p* < 0.001; Supplementary Table S3). The concentration of cortisol in maternal saliva decreased as the season progressed (Figure 3g) and tended to be higher in 2020, the year of higher food availability (Figure 3h).

## Discussion

We used a fully crossed, repeated measures design to characterize individual variation in baseline cortisol levels in a wild population of Antarctic fur seals. We found that baseline cortisol was only marginally explained by genetic factors, while high within-individual repeatability and among-individual variability in both pups and mothers could be largely explained by a combination of intrinsic and extrinsic factors. Our results provide detailed insights into the individualization of an endocrine phenotype in a wild pinniped population.

### Heritability estimates

We quantified the narrow sense heritability of baseline cortisol using animal models based on a simple pedigree and a SNP array. The former approach assumes that individuals of unknown parentage are unrelated to all other individuals in the population, which can lead to heritability being underestimated (Kruuk 2004). By contrast, genomic approaches are capable of quantifying unbiased relatedness for all sampled individuals, but can be time-consuming and costly to produce (Frentiu et al. 2008). Despite these differences, we found that both approaches produced consistently low heritability estimates for baseline cortisol in Antarctic fur seals. Heritability can be low because of low additive genetic variation, high environmental variance, or cross-environment genetic correlations (i.e. the genetic basis of the trait varies between different environments) (Charmantier and Garant 2005). Our low estimates, which stand in contrast to our original expectations and previously published empirical estimates from other species (Jenkins et al. 2014; Stedman et al. 2017; Bairos-Novak et al. 2018), might therefore be reflective of the extreme heterogeneity of the environmental conditions encountered by the study population at Bird Island. This explanation would be in line with other empirical studies of wild vertebrate populations showing a decrease in heritability under poor environmental conditions (e.g. (Wilson et al. 2006)). Alternatively, low heritability may reflect the different life stages at which cortisol concentrations were measured (i.e. mothers *verses* pups).

### Intrinsic and extrinsic factors affecting baseline cortisol

For both pups and mothers, we were able to explain a substantial proportion of the total phenotypic variance in baseline cortisol by including individual-based and environmental variables in our models (conditional *R*^2^ = 0.66 for pups and *R*^2^ = 0.77 for mothers). Similar results have been obtained for a variety of species (e.g. (Joly and Cameron 2018; Uchida et al. 2021), but see (Azevedo et al. 2019)), suggesting that contextual and developmental phenotypic plasticity in response to environmental heterogeneity may be a widespread phenomenon. In addition, we found that baseline cortisol levels were consistent within individuals, suggesting that individualized endocrine phenotypes become established during early ontogeny and persist at least until nutritional independence. Baseline cortisol might therefore represent a stable attribute by which fur seals adapt to spatial or temporal heterogeneity in their environment (Réale and Dingemanse 2010).

Our models uncovered a strong influence of age and days postpartum on baseline cortisol levels in pups and mothers, respectively, with salivary cortisol decreasing over time. One explanation for this may be the shifting environmental conditions that individuals experience as the season progresses. Pregnant females arrive ashore in December and give birth on crowded breeding beaches. Mothers continue to suckle their pups on the beach until about 30 days postpartum, when most females transition into the more sheltered and less crowded tussock grass that covers most of the island’s interior (Doidge et al. 1984). Also during this time, adult males begin to abandon their territories and migrate to higher latitudes around the Antarctic ice shelf (Forcada and Staniland 2018). Consequently, the frequency of unpredictable challenges for both pups and mothers likely declines as the season progresses. A corresponding reduction in baseline cortisol is therefore in line with previous research suggesting that cortisol levels are lower under stable environmental conditions compared with environments associated with frequent disturbances (Fairbanks et al. 2011).

We detected significantly higher cortisol concentrations among pups and mothers sampled in 2020, the year of higher food availability. This was surprising given the many empirical studies that have linked elevated cortisol concentrations to food shortages and periods of nutritional stress (e.g. (Kitaysky et al. 2007; Behie et al. 2010; Bryan et al. 2013; Garber et al. 2020)). We can think of two possible explanations for this pattern. On the one hand, our results could be explained by higher population densities in the second year of our study, as significantly more females bred in 2020 compared to 2019 (Nagel et al. 2021). This would be in line with the small, albeit non-significant, effect of colony on baseline cortisol, with hormone concentrations being marginally higher in both pups and mothers at SSB compared to FWB. On the other hand, circulating levels of cortisol are essential for the maintenance of metabolic functions (Kuo et al. 2015), and food-induced cortisol secretions have been documented in the literature (Quigley and Yen 1979; Gibson et al. 1999; Stimson et al. 2014). Shorter foraging trip durations (Nagel et al. 2021) and consequently more frequent meals for pups and mothers in 2020 may have resulted in higher average baseline cortisol concentrations, which facilitate protein and carbohydrate metabolism.

More indicative of the hypothesized correlation between cortisol and nutritional stress, we found a significant negative relationship between baseline cortisol levels and weight in pups. Elevated cortisol levels provide individuals with a source of energy by stimulating gluconeogenesis, which increases the delivery of glucose into the bloodstream (Wingfield and Romero 2011). Furthermore, cortisol can enhance fat oxidation by other mechanisms, like promoting production of hormone sensitive lipase (Samra et al. 1996). Given that pups must tolerate bouts of fasting lasting up to 11 days while their mothers forage at sea (Forcada and Staniland 2018), our results may reflect a physiological response to prolonged periods of natural food limitation (Ortiz et al. 2001; Jeanniard du Dot et al. 2009). In other words, fasting pups may increase baseline cortisol to release energy, resulting in a reduction of absolute body fat and overall weight. This would also explain why we do not see a similar relationship in mothers, who remain ashore between foraging trips for as little as 24 hours and on average only two days (Boyd et al. 1991). Alternatively, pups may be more susceptible to environmental stressors than adult females, with lighter pups requiring more energy to maintain homeostasis under, for example, unfavorable climatic conditions.

In pups, we also uncovered a significant association between baseline cortisol and sex, with hormone concentrations being moderately higher in males than females. Previous studies of the sex-specific secretion of cortisol have produced diverse results, including conflicting evidence from within a single species (Steller sea lion pups: (Keogh et al. 2010; Myers et al. 2010). These contrasting findings highlight the complexity of interactions between cortisol and the sex hormones, which can vary with the reproductive system, phase, and cycle (Levine 2002). In addition, the social and environmental stressors associated with growth and reproduction in each sex are likely to vary among species. For example, male Antarctic fur seal pups engage more often in social interactions and risk prone behaviors (Jones et al. 2020) such that the number of “stressful” events encountered may be higher in males than females.

Contrary to our initial expectations, colony only had a marginal, non-significant effect on baseline salivary cortisol in pups and their mothers. This is in contrast to a previous study where cortisol concentrations, measured from hair, were higher in mothers (but not offspring) from the high density colony (Meise et al. 2016). We can think of two non-mutually exclusive explanations for this discrepancy. First, hair cortisol concentrations are thought to reflect events in the recent past (hours or days) (Kalliokoski et al. 2019), whereas salivary cortisol captures circulating hormone levels at the immediate time of sampling (Lewis 2006). Consequently, hair concentrations of baseline cortisol might integrate a larger number of stressful events, allowing density-dependent differences to more readily accumulate. Second, the first study was conducted in 2011 when population densities were much higher at SSB (approximately 568 breeding females were observed at SSB in 2011 compared with 282 in 2019 and 409 in 2020, a reduction of 50% and 28%, respectively (Forcada and Hoffman 2014; Nagel et al. 2021), which may have accentuated differences between the two colonies.

### Strengths, limitations, and future directions

The past decade has witnessed a shift in our understanding of individual differences from a perceived statistical nuisance to a fundamental and compelling aspect of behavioral ecology and evolutionary biology (Bolnick et al. 2003; Trillmich et al. 2018; Krüger et al. 2021). Our study contributes towards this narrative by decomposing individual variation in endocrine phenotypes across different life history stages and environments in Antarctic fur seal pups and their mothers. Our results are indicative of substantial developmental plasticity in the HPA axis of pups, with individualized endocrine phenotypes becoming established during early ontogeny and persisting at least until nutritional independence. Baseline cortisol may therefore help to facilitate the match between an individual’s phenotype and the environment. That we find a higher relative contribution of the environment compared to genetic factors to phenotypic variation further suggests that the endocrine phenotype may be highly adaptable to unpredictable environmental conditions. This has implications for our understanding of population dynamics, both in the declining Antarctic fur seal population (Forcada and Hoffman 2014) and other in systems impacted by climate change.

However, further research is needed to elucidate both the fitness consequences of endocrine variability and how this may respond to environmental heterogeneity over longer timescales. To meet the strictest definition of phenotypic plasticity, individual variation in the focal trait must be demonstrated in different environments, ideally across multiple years (Boutin and Lane 2014). For developmental plasticity, an individual’s phenotype must persist into later life stages, while changing reaction norms over an individual’s lifetime would suggest contextual plasticity (Nettle and Bateson 2015; Trillmich et al. 2015). Our conclusions are also limited by the comparison of only two colonies across two seasons. While including additional colonies in the study would be logistically prohibitive given the inaccessibility of most breeding beaches, a continuation of this study across additional breeding seasons would certainly be feasible. Increasing the duration of this study would be particularly relevant given the ever increasing occurrence and intensity of severe weather events in the sub-Antarctic region (Turner et al. 2005).

Overall, our work builds upon the existing literature on the heritability and phenotypic plasticity of baseline cortisol in response to environmental heterogeneity. Although our results are framed in the context of the Antarctic fur seal, our study has implications for understanding the importance of endocrine mechanisms in other populations. We can contribute towards a growing body of evidence showing that endocrine phenotypes are ecologically important traits that can potentially affect population dynamics through their influence on life history traits. Expanding our understanding of those extrinsic and intrinsic factors that influence baseline cortisol therefore gives insight into the factors that potentially limit or improve population persistence in a changing environment.

## Supporting information

R Markdown

## Acknowledgements

The authors would like to thank Ana Bertoldi Carneiro, Freya Blockley, Jamie Coleman, Alexandra Dodds, Vicki Foster, Derren Fox, Iain Angus Gordon, Pauline Goulet, Rosie Hall, Cary Jackson, Adam Lowndes, Elizabeth Morgan, Rachael Orben, Jessica Ann Philips, David Reid, and Mark Whiffin for additional help in the field. We kindly thank Sabine Kruse for conducting the endocrine analyses. We are also grateful to Jonas Schwartz for tips on how to collect saliva samples from young pups in the field. A special thanks to Océane Salles for producing the fur seal cartoon in Figure 1c.

## Author contributions

RN, CS, CT, and CF-C collected the data. SK advised on field protocols and coordinated cortisol quantification. RN analyzed the data. AJP generated the SNP genotype dataset and DLJV built the genomic relatedness matrix. RN and CC performed the molecular laboratory work. JIH, SK, and JF conceived and developed the project. RN drafted the manuscript. All of the authors commented on and approved the final manuscript.

## Data Availability Statement

Our code and accompanying documentation are provided in the form of an R Markdown file (Supplementary Information). The data used for this study are available via Zenodo, https://doi.org/10.5281/zenodo.6336716.

