## Supplementary material for "Low heritability and high phenotypic plasticity of salivary cortisol in response to environmental heterogeneity in a wild pinniped": R Markdown

### R-code for 'Saliva cortisol - Antarctic fur seals'

compiled by XXX

**Manuscript figures** **28**

---

This document provides all the `R code` for the manuscript titled *Low heritability and high phenotypic plasticity of cortisol in response to environmental heterogeneity in fur seals* by XXX. Both the R Markdown file and the raw data can be downloaded via DRYAD (XXXX). If you have any questions, don't hesitate to contact XXX.

The data was collected from Antarctic fur seal (*Arctocephalus gazella*) mother-pup pairs on Bird Island, South Georgia between 2018-2020.

---

#### Download packages and libraries

In order to repeat analyses presented in this manuscript a number of packages that extend the functionalities of base `R` are required. These can be installed using the code `install.packages("xxPACKAGENAMExx", dependencies = TRUE)`

```
library(tidyverse)
library(reshape2)
library(data.table)
library(plyr)
library(gdata)
library(ggpubr)
library(ggforce)
library(patchwork)
library(lme4)
library(mgcv)
library(goft)
library(DHARMa)
library(performance)
library(MuMIn)
library(specr)
library(car)
library(emmeans)
library(MCMCglmm)
set.seed(529843)
```

---

#### Data

##### Salivary cortisol data

```
# read in data
cortisol <- read.csv("cortisol_2018-2020_AllData.csv", header = TRUE, sep = ",")

# fix column categories
cortisol$Season <- as.character(cortisol$Season)

# clean up dataset
cortisol <- subset(cortisol, select = -c(Sample_Label, RohWert1, RohWert2, Verd.Faktor,
    EndWert1, EndWert2, Aggressiveness, LocalDensity, Length_Avg))

col_order <- c("ID", "ID_status", "ID_pair", "Season", "Beach", "Date", "Day_Actual",
    "Day", "CatchTime", "Sex", "Weight", "Girth", "Span", "Length", "CI", "Death",
    "Cause_of_death", "SampleCollected", "Time", "TimeBetweenSamples",
    "Mittelwert_ng.ml", "Bemerkung")
cortisol <- cortisol[, col_order]
rm(col_order)

# remove nas
```

```
cortisol.rmna <- cortisol[!is.na(cortisol$Mittelwert_ng.ml), ]

# visually inspect data
plot(cortisol.rmna$Mittelwert_ng.ml, ylab = "cortisol concentration (ng/mL)", pch = 20)
```

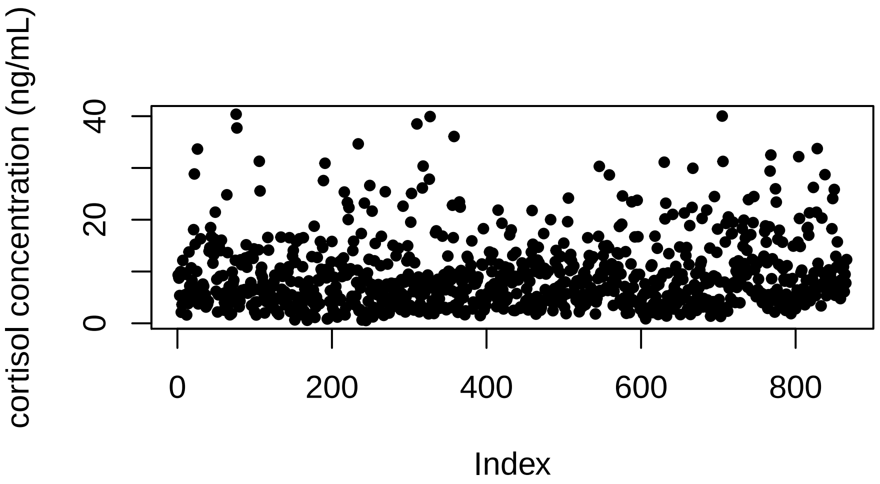

```
hist(cortisol.rmna$Mittelwert_ng.ml)
```

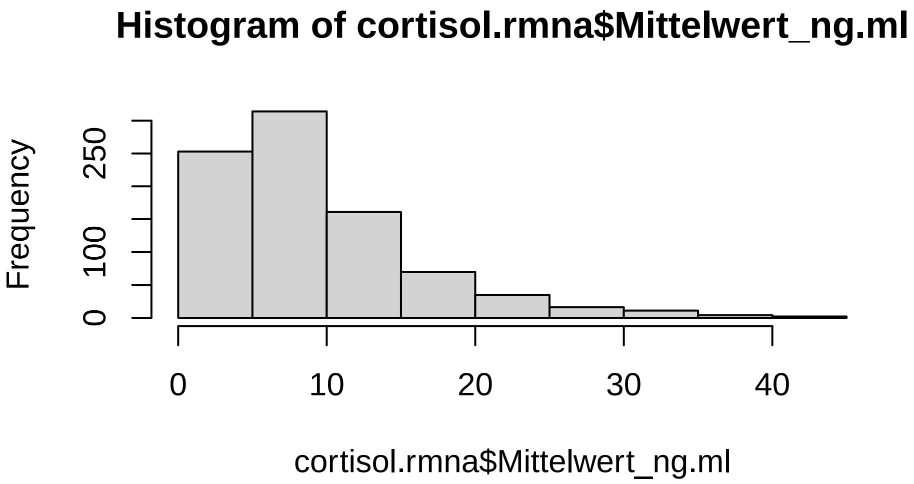

##### Pedigree and relatedness matrix data

```
#### SNP relatedness matrix ----
snp <- read.table("AFS_grm.txt", header = TRUE, sep = "\t")
```

```
# over-write bas id with study id
IDs <- read.csv("Mum-Pup_BASid.csv", header = TRUE, sep = ",")

names(IDs)[names(IDs) == "BAS_ID"] <- "Sample_1"
snp <- left_join(snp, IDs, by = "Sample_1")
snp <- data.frame(snp[, -1])
names(snp)[names(snp) == "ID"] <- "Sample_1"

names(IDs)[names(IDs) == "Sample_1"] <- "Sample_2"
snp <- left_join(snp, IDs, by = "Sample_2")
snp <- data.frame(snp[, -1])
names(snp)[names(snp) == "ID"] <- "Sample_2"

# make list into a matrix
snp.matrix <- reshape(snp, idvar = "Sample_1", timevar = "Sample_2", v.names = "rxy",
    direction = "wide")
names(snp.matrix) <- substring(names(snp.matrix), 5)
snp.matrix <- data.frame(snp.matrix[, -1], row.names = snp.matrix[, 1])

# fill matrix on both sides
snp.matrix <- t(snp.matrix)
lowerTriangle(snp.matrix) = upperTriangle(snp.matrix, byrow = TRUE)
```

### Cortisol baseline

#### Pups

To determine whether weight, body condition, age, sex, death, season, colony and an interaction between season and colony explained a significant proportion of the variation in baseline cortisol among pups, we fit a generalized linear mixed model (GLMER).

```
# trim dataset down & remove NAs
cortisol.pup.baseline <- cortisol[cortisol$ID_status == "pup" & cortisol$SampleCollected
    == "Baseline", ]
cortisol.pup.baseline <- cortisol.pup.baseline[complete.cases(cortisol.pup.baseline[,
    c("Mittelwert_ng.ml", "Day_Actual", "Season", "Beach", "Sex", "CI")]), ]
```

##### Multicollinearity

Prior to building the model, we needed to account for both structural multicollinearity (body condition is calculated in part from pup weight) and data multicollinearity (pup age and weight are highly correlated (R = 0.8)). Body condition, weight and age were centered by subtracting the mean from all observed values.

```
## weight and age are not independent variables (data multicollinearity)
ggscatter(cortisol.pup.baseline, x = "Day_Actual", y = "Weight", add = "reg.line",
    conf.int = TRUE, cor.coef = TRUE, cor.method = "pearson", na.rm = TRUE)
```

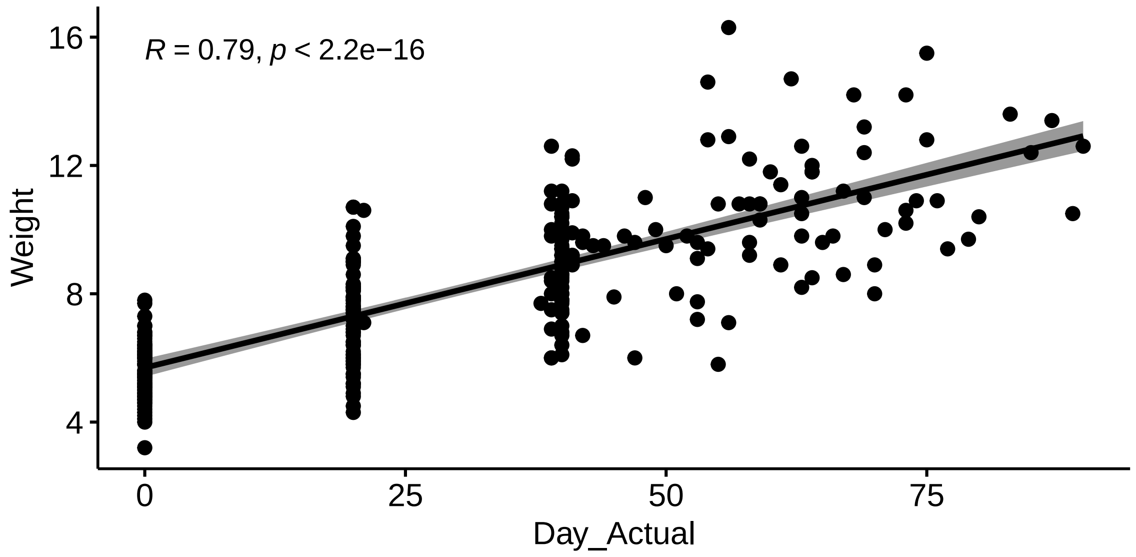

```
## weight and ci are not independent variables (structural multicollinearity)
ggscatter(cortisol.pup.baseline, x = "Weight", y = "CI", add = "reg.line", conf.int =
    TRUE, cor.coef = TRUE, cor.method = "pearson", na.rm = TRUE)
```

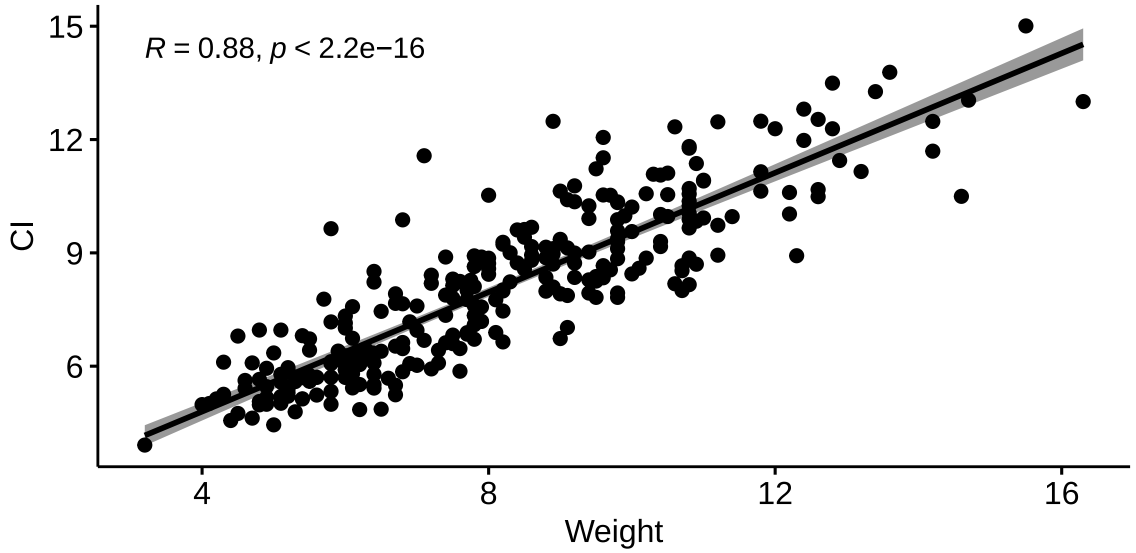

```
## --> taken into account by rescaling and centering these continuous parameters
## Center the variables by subtracting the mean. i.e. calculated mean for each
## continuous independent variable is subtracted from all observed values of that
## variable
cortisol.pup.baseline.rsc <- cortisol.pup.baseline
cortisol.pup.baseline.rsc[, c("Day", "Day_Actual", "CI", "Weight")] <-
    scale(cortisol.pup.baseline.rsc[, c("Day", "Day_Actual", "CI", "Weight")])
```

##### Preliminary analyses

We specified the a gamma conditional distribution of the response variable (pup baseline cortisol) with a log link function.

In preliminary analyses, we tested (1) if individual variation in baseline cortisol was repeatable across measurements by allowing the linear regression fit for each individual pup to differ in intercept and (2) for the presence of heterogeneous variance by allowing individual slopes to vary by age. Including pup ID as a random intercept and age as a random slope significantly improved model fit.

```
#### error distribution ---- look at distribution of response variable
hist(cortisol.pup.baseline.rsc$Mittelwert_ng.ml, breaks = 100)
```

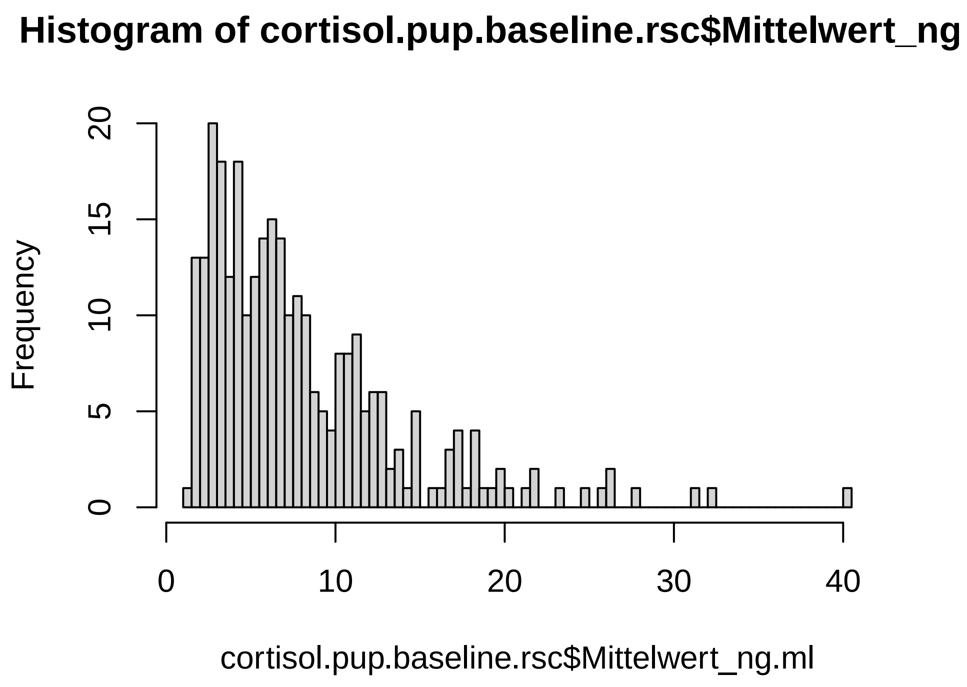

```r
# null hypothesis = distribution follows gamma / inverse gamma distribution
goft::gamma_test(cortisol.pup.baseline.rsc$Mittelwert_ng.ml)
```

```
##
##  Test of fit for the Gamma distribution
##
## data:  cortisol.pup.baseline.rsc$Mittelwert_ng.ml
## V = 4.1784, p-value = 0.003131
```

```r
goft::ig_test(cortisol.pup.baseline.rsc$Mittelwert_ng.ml)
```

```
## [[1]]
##
##  Test for Inverse Gaussian distributions using a transformation to
##  normality
##
## data:  cortisol.pup.baseline.rsc$Mittelwert_ng.ml
## W = 0.99458, p-value = 0.3952
##
##
## [[2]]
##
##  Test for Inverse Gaussian distributions using a transformation to
##  gamma variables
##
## data:  cortisol.pup.baseline.rsc$Mittelwert_ng.ml
## AD = 0.44974, p-value = 0.6332
## alternative hypothesis: cortisol.pup.baseline.rsc$Mittelwert_ng.ml  does not follow an Inverse Gauss
```

```
#### run glmer ---- inverse gamma error distribution
m1 <- glmer(Mittelwert_ng.ml ~ Day_Actual + CI + Weight + Season * Beach + Sex + (1 |
    ID), family = Gamma(link = "log"), data = cortisol.pup.baseline.rsc)

# 1. behavioral repeatable? i.e. better model fit when including ID as random
# effect?
m1.noID <- glm(Mittelwert_ng.ml ~ Day_Actual + CI + Weight + Season * Beach + Sex, family
    = Gamma(link = "log"), data = cortisol.pup.baseline.rsc)

# 2. fit glmer with random slopes and intercepts. i.e. individual slopes can vary
# depending on age = strength of the effect (slope) of age on cortisol varies
# depending on individual (see Barr et al. (2013) and Harrison et al. (2018))
m2 <- glmer(Mittelwert_ng.ml ~ Day_Actual + CI + Weight + Season * Beach + Sex +
    (Day_Actual | ID), family = Gamma(link = "log"), data = cortisol.pup.baseline.rsc)
#### if the model doesn't converge --> re-run with more iterations
ss <- getME(m2, c("theta", "fixef"))
m2 <- update(m2, start = ss, control = glmerControl(optimizer = "bobyqa", optCtrl =
    list(maxfun = 2e+05)))

# best model fit with random intercept (ID) and random slope (age)
anova(m1, m1.noID, m2)
```

```
## Data: cortisol.pup.baseline.rsc
## Models:
## m1.noID: Mittelwert_ng.ml ~ Day_Actual + CI + Weight + Season * Beach + Sex
## m1: Mittelwert_ng.ml ~ Day_Actual + CI + Weight + Season * Beach + Sex + (1 | ID)
## m2: Mittelwert_ng.ml ~ Day_Actual + CI + Weight + Season * Beach + Sex + (Day_Actual | ID)
##         npar    AIC    BIC  logLik deviance  Chisq Df Pr(>Chisq)
## m1.noID    9 1556.8 1589.9 -769.41   1538.8
## m1        10 1543.0 1579.7 -761.49   1523.0 15.842  1  6.886e-05 ***
## m2        12 1488.9 1532.9 -732.43   1464.9 58.130  2  2.383e-13 ***
## ---
## Signif. codes:  0 '***' 0.001 '**' 0.01 '*' 0.05 '.' 0.1 ' ' 1
```

##### Residual check of full model

The residuals of the full model were visually inspected for linearity and equality of error variances (using plots of residuals versus fits), normality (using Q–Q plots) and homogeneity of variance (using Levene's Test).

```
# check for singularity = some of the constrained parameters of the random
# effects theta parameters are on the boundary (equal to or very close to zero)
isSingular(m2, tol = 1e-04)
```

```
## [1] FALSE
```

```
# Check for independence & homoscedasticity - points should be randomly scattered
# around zero for the entire range of fitted values
plot(fitted(m2), resid(m2, type = "pearson"), main = "Residuals vs Fitted")
abline(0, 0, col = "red")
```

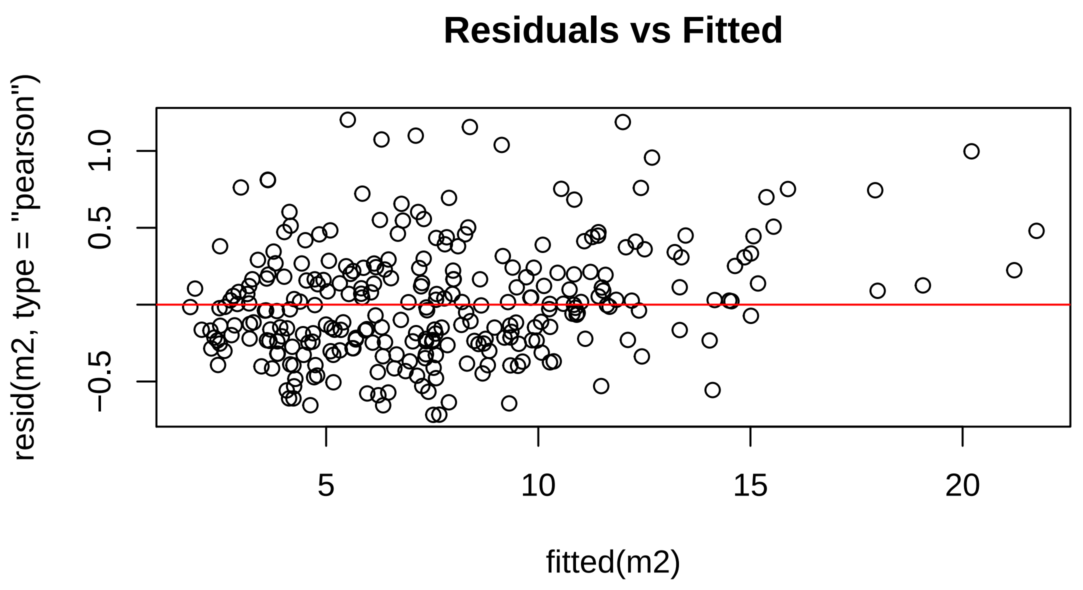

```
# Check if residuals are normally distributed - points should follow the line.
qqnorm(resid(m2), main = "Normal Q-Q")
qqline(resid(m2), col = "red")
```

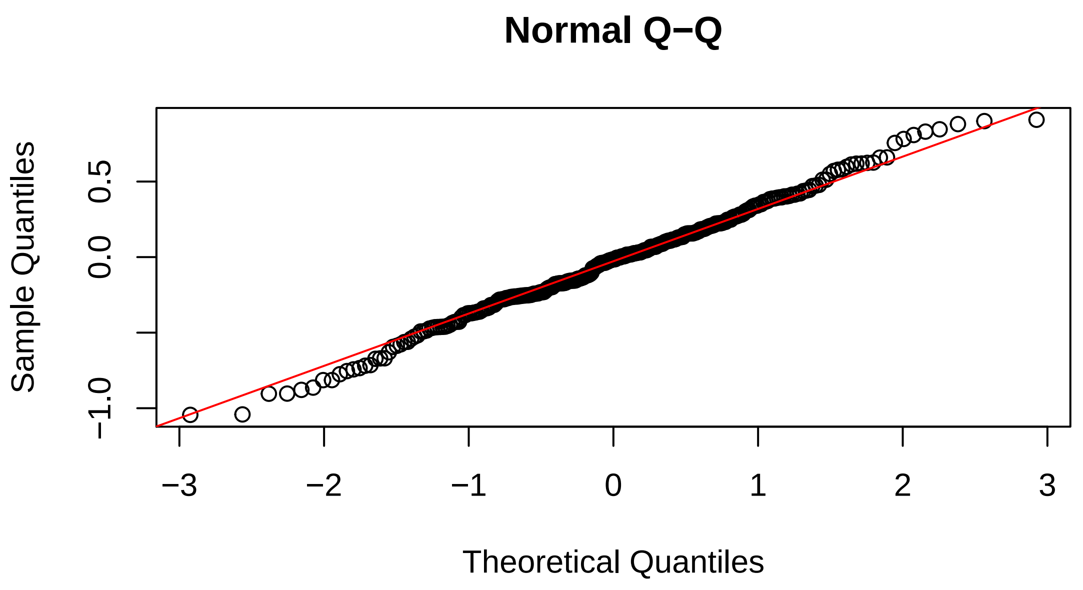

```
# Additional check for homogeneity of variance - Levene's Test: 1. Extract
# residuals and place them in a new column in the original data table
cortisol.pup.baseline.rsc$m2.Res <- residuals(m2)
# 2. Create a new column with the absolute value of the residuals
cortisol.pup.baseline.rsc$Abs.m2.Res <- abs(cortisol.pup.baseline.rsc$m2.Res)
```

```r
# 3. Square the absolute values of the residuals to provide more robust estimate
cortisol.pup.baseline.rsc$m2.Res2 <- cortisol.pup.baseline.rsc$Abs.m2.Res^2
# 4. ANOVA of the squared residuals. A p > 0.05 means that the variance of the
# residuals is equal and the assumption of homoscedasticity is met
Levene.Model.F <- lm(m2.Res2 ~ ID, data = cortisol.pup.baseline.rsc)
anova(Levene.Model.F)
```

```
## Analysis of Variance Table
##
## Response: m2.Res2
##            Df Sum Sq  Mean Sq F value  Pr(>F)
## ID         95 4.3933 0.046246  1.2842 0.07368 .
## Residuals 194 6.9861 0.036011
## ---
## Signif. codes:  0 '***' 0.001 '**' 0.01 '*' 0.05 '.' 0.1 ' ' 1
```

##### Model reduction

A backward elimination based on the chi-squared statistic was implemented to simplify the model. The best fit model was determined based on AIC values. The statistical significance of fixed predictors was assessed using a Wald test.

```r
#### model reduction, backwards elimination
Anova(m2)
```

```
## Analysis of Deviance Table (Type II Wald chisquare tests)
##
## Response: Mittelwert_ng.ml
##                  Chisq Df Pr(>Chisq)
## Day_Actual     14.2407  1  0.0001609 ***
## CI              2.8788  1  0.0897519 .
## Weight         18.9316  1  1.355e-05 ***
## Season          9.2111  1  0.0024056 **
## Beach           2.6743  1  0.1019811
## Sex             8.4943  1  0.0035626 **
## Season:Beach    1.3978  1  0.2370957
## ---
## Signif. codes:  0 '***' 0.001 '**' 0.01 '*' 0.05 '.' 0.1 ' ' 1
```

```r
m3 <- update(m2, . ~ . - Season:Beach)
Anova(m3)
```

```
## Analysis of Deviance Table (Type II Wald chisquare tests)
##
## Response: Mittelwert_ng.ml
##              Chisq Df Pr(>Chisq)
## Day_Actual 13.6781  1   0.000217 ***
## CI          2.8279  1   0.092639 .
## Weight     19.7132  1  8.998e-06 ***
## Season      8.9942  1   0.002708 **
## Beach       2.6003  1   0.106843
```

```
## Sex         8.1161  1   0.004387 **
## ---
## Signif. codes:  0 '***' 0.001 '**' 0.01 '*' 0.05 '.' 0.1 ' ' 1
```

```r
m4 <- update(m3, . ~ . - Beach)
Anova(m4)
```

```
## Analysis of Deviance Table (Type II Wald chisquare tests)
##
## Response: Mittelwert_ng.ml
##               Chisq Df Pr(>Chisq)
## Day_Actual 13.9424  1  0.0001885 ***
## CI          2.8142  1  0.0934338 .
## Weight     18.1750  1  2.015e-05 ***
## Season      8.5768  1  0.0034047 **
## Sex         6.2794  1  0.0122151 *
## ---
## Signif. codes:  0 '***' 0.001 '**' 0.01 '*' 0.05 '.' 0.1 ' ' 1
```

```r
m5 <- update(m4, . ~ . - CI)
Anova(m5)
```

```
## Analysis of Deviance Table (Type II Wald chisquare tests)
##
## Response: Mittelwert_ng.ml
##               Chisq Df Pr(>Chisq)
## Day_Actual 11.1901  1  0.0008223 ***
## Weight     17.0072  1  3.724e-05 ***
## Season      9.3178  1  0.0022694 **
## Sex         5.3926  1  0.0202219 *
## ---
## Signif. codes:  0 '***' 0.001 '**' 0.01 '*' 0.05 '.' 0.1 ' ' 1
```

```r
m6 <- update(m5, . ~ . - Sex)
Anova(m6)
```

```
## Analysis of Deviance Table (Type II Wald chisquare tests)
##
## Response: Mittelwert_ng.ml
##               Chisq Df Pr(>Chisq)
## Day_Actual 13.5050  1  0.0002379 ***
## Weight     12.8245  1  0.0003421 ***
## Season      8.3994  1  0.0037534 **
## ---
## Signif. codes:  0 '***' 0.001 '**' 0.01 '*' 0.05 '.' 0.1 ' ' 1
```

```r
m7 <- update(m6, . ~ . - Season)
Anova(m7)
```

```
## Analysis of Deviance Table (Type II Wald chisquare tests)
##
```

```
## Response: Mittelwert_ng.ml
##             Chisq Df Pr(>Chisq)
## Day_Actual 12.957  1  0.0003188 ***
## Weight     11.838  1  0.0005804 ***
## ---
## Signif. codes:  0 '***' 0.001 '**' 0.01 '*' 0.05 '.' 0.1 ' ' 1
```

```r
anova(m2, m3, m4, m5, m6, m7)
```

```
## Data: cortisol.pup.baseline.rsc
## Models:
## m7: Mittelwert_ng.ml ~ Day_Actual + Weight + (Day_Actual | ID)
## m6: Mittelwert_ng.ml ~ Day_Actual + Weight + Season + (Day_Actual | ID)
## m5: Mittelwert_ng.ml ~ Day_Actual + Weight + Season + Sex + (Day_Actual | ID)
## m4: Mittelwert_ng.ml ~ Day_Actual + CI + Weight + Season + Sex + (Day_Actual | ID)
## m3: Mittelwert_ng.ml ~ Day_Actual + CI + Weight + Season + Beach + Sex + (Day_Actual | ID)
## m2: Mittelwert_ng.ml ~ Day_Actual + CI + Weight + Season * Beach + Sex + (Day_Actual | ID)
##    npar    AIC    BIC  logLik deviance  Chisq Df Pr(>Chisq)
## m7    7 1498.6 1524.3 -742.31   1484.6
## m6    8 1492.8 1522.1 -738.39   1476.8 7.8364  1   0.005121 **
## m5    9 1489.6 1522.6 -735.78   1471.6 5.2286  1   0.022218 *
## m4   10 1488.8 1525.5 -734.37   1468.8 2.8069  1   0.093859 .
## m3   11 1488.2 1528.6 -733.12   1466.2 2.5065  1   0.113377
## m2   12 1488.9 1532.9 -732.43   1464.9 1.3853  1   0.239201
## ---
## Signif. codes:  0 '***' 0.001 '**' 0.01 '*' 0.05 '.' 0.1 ' ' 1
```

```r
summary(m3)
```

```
## Generalized linear mixed model fit by maximum likelihood (Laplace
##   Approximation) [glmerMod]
##  Family: Gamma  ( log )
## Formula: Mittelwert_ng.ml ~ Day_Actual + CI + Weight + Season + Beach +
##     Sex + (Day_Actual | ID)
##    Data: cortisol.pup.baseline.rsc
## Control: glmerControl(optimizer = "bobyqa", optCtrl = list(maxfun = 2e+05))
##
##      AIC      BIC   logLik deviance df.resid
##   1488.2   1528.6   -733.1   1466.2      279
##
## Scaled residuals:
##      Min       1Q   Median       3Q      Max
## -1.66923 -0.56407 -0.03717  0.52412  2.91354
##
## Random effects:
##  Groups   Name        Variance Std.Dev. Corr
##  ID       (Intercept) 0.05081  0.2254
##           Day_Actual  0.06466  0.2543   0.19
##  Residual             0.18237  0.4270
## Number of obs: 290, groups:  ID, 96
##
## Fixed effects:
##             Estimate Std. Error t value Pr(>|z|)
```

```
## (Intercept)  1.57547    0.08878  17.745  < 2e-16 ***
## Day_Actual  -0.28911    0.07817  -3.698 0.000217 ***
## CI           0.14287    0.08496   1.682 0.092639 .
## Weight      -0.36126    0.08136  -4.440    9e-06 ***
## Season1920   0.24990    0.08333   2.999 0.002708 **
## BeachSSB     0.13266    0.08227   1.613 0.106843
## SexM         0.24435    0.08577   2.849 0.004387 **
## ---
## Signif. codes:  0 '***' 0.001 '**' 0.01 '*' 0.05 '.' 0.1 ' ' 1
##
## Correlation of Fixed Effects:
##            (Intr) Dy_Act CI     Weight Ss1920 BchSSB
## Day_Actual -0.032
## CI         -0.043 -0.485
## Weight      0.192 -0.204 -0.616
## Season1920 -0.452  0.065 -0.070  0.027
## BeachSSB   -0.581  0.037  0.001 -0.090 -0.011
## SexM       -0.601  0.094  0.115 -0.279  0.007  0.207
```

```
# Check model for Multicollinearity
performance::check_collinearity(m3)
```

```
## # Check for Multicollinearity
##
## Low Correlation
##
##   Parameter  VIF Increased SE
##  Day_Actual 2.81         1.68
##          CI 4.37         2.09
##      Weight 3.63         1.90
##      Season 1.01         1.00
##       Beach 1.05         1.03
##         Sex 1.13         1.06
```

```
# variance estimates: variance of ID (Intercept) = variability of the intercept
# across ID & variance of ID (day) = variability in the slope across ID &
# residual variance = s2 = residual variation around the regression line
vc <- lme4::VarCorr(m3)
print(vc, comp = c("Variance", "Std.Dev."), digits = 2)
```

```
##  Groups   Name        Variance Std.Dev. Corr
##  ID       (Intercept) 0.051    0.23
##           Day_Actual  0.065    0.25     0.19
##  Residual             0.182    0.43
```

```
# Marginal and conditional R2
r.squaredGLMM(m3)
```

```
##                 R2m       R2c
## delta     0.4395988 0.6566150
## lognormal 0.4522431 0.6755013
## trigamma  0.4254723 0.6355147
```

```r
# ICC = percent of the variation in outcome attributable to differences between
# random effect
icc_specs(m3, percent = TRUE)
```

```
##        grp       vcov        icc   percent
## 1       ID 0.05081243 0.16468770 16.468770
## 2       ID 0.06466452 0.20958360 20.958360
## 3       ID 0.01069463 0.03466226  3.466226
## 4 Residual 0.18236652 0.59106644 59.106644
```

---

#### Mums

To determine whether weight, body condition, time during the season (i.e. just after birth or just before weaning her pup), season, colony and an interaction between season and colony explained a significant proportion of the variation in baseline cortisol among mothers, we fit a GLMER.

```r
# trim dataset down & remove NAs
cortisol.mum.baseline <- cortisol[cortisol$ID_status == "mum" & cortisol$SampleCollected
    == "Baseline", ]
cortisol.mum.baseline <- cortisol.mum.baseline[complete.cases(cortisol.mum.baseline[,
    c("Mittelwert_ng.ml", "Day_Actual", "Season", "Beach", "CI")]), ]
```

##### Multicollinearity

Prior to building the model, we needed to account for both structural multicollinearity (body condition is calculated in part from weight) and data multicollinearity (time during the season and weight are highly correlated). Body condition, weight and age were centered by subtracting the mean from all observed values.

```r
# weight, age and ci are not independent variables - taken into account by
# rescaling and centering these continuous parameters
cortisol.mum.baseline.rsc <- cortisol.mum.baseline

cortisol.mum.baseline.rsc[, c("Day", "Day_Actual", "CI", "Weight")] <-
    scale(cortisol.mum.baseline.rsc[, c("Day", "Day_Actual", "CI", "Weight")])
```

##### Preliminary analyses

We specified a gamma conditional distribution of the response variable (maternal baseline cortisol) with a log link function.

Testing for the presence of heterogeneous variance by allowing individual slopes to vary by time during the season created a random effects structure that was too complex to be supported by data. Including maternal ID as a random intercept significantly improved model fit.

```r
#### error distribution ---- look at distribution of response variable
hist(cortisol.mum.baseline.rsc$Mittelwert_ng.ml, breaks = 100)
```

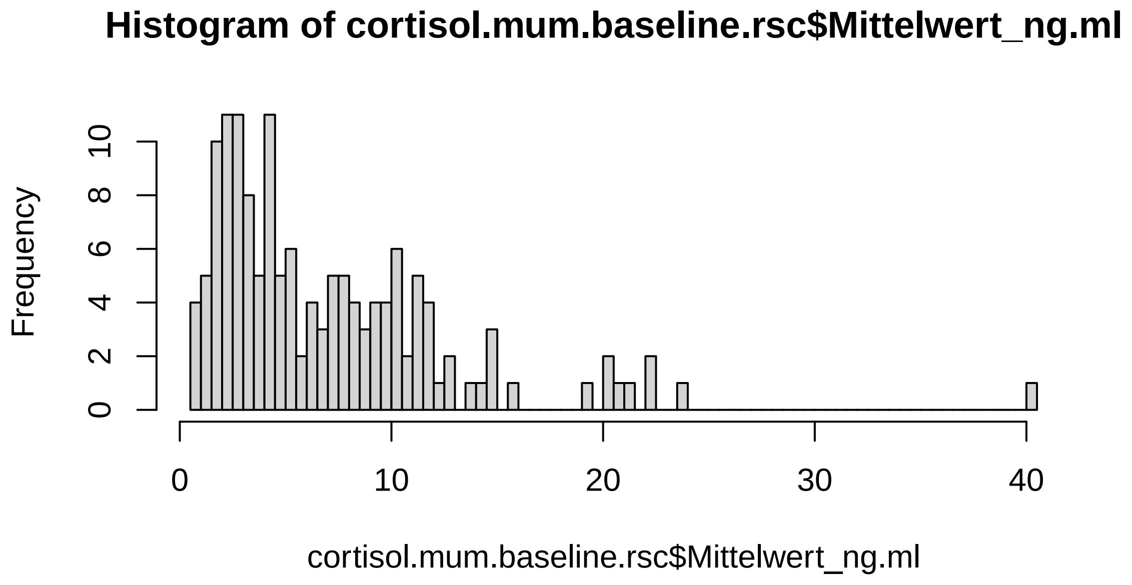

```r
# null hypothesis = distribution follows gamma / inverse gamma distribution
goft::gamma_test(cortisol.mum.baseline.rsc$Mittelwert_ng.ml)
```

```
##
##  Test of fit for the Gamma distribution
##
## data:  cortisol.mum.baseline.rsc$Mittelwert_ng.ml
## V = 2.1757, p-value = 0.1239
```

```r
goft::ig_test(cortisol.mum.baseline.rsc$Mittelwert_ng.ml)
```

```
## [[1]]
##
##  Test for Inverse Gaussian distributions using a transformation to
##  normality
##
## data:  cortisol.mum.baseline.rsc$Mittelwert_ng.ml
## W = 0.9879, p-value = 0.2395
##
##
## [[2]]
##
##  Test for Inverse Gaussian distributions using a transformation to
##  gamma variables
##
## data:  cortisol.mum.baseline.rsc$Mittelwert_ng.ml
## AD = 0.51625, p-value = 0.545
## alternative hypothesis: cortisol.mum.baseline.rsc$Mittelwert_ng.ml  does not follow an Inverse Gauss[illegible]
```

```r
#### GLMER ----
mm1.log <- glmer(Mittelwert_ng.ml ~ Day_Actual + CI + Weight + Season * Beach + (1 | ID),
    family = Gamma(link = "log"), data = cortisol.mum.baseline.rsc)

# behavioral repeatable?
mm1.noID <- glm(Mittelwert_ng.ml ~ Day_Actual + CI + Weight + Season * Beach, family =
    Gamma(link = "log"), data = cortisol.mum.baseline.rsc)

# add random slope? mm2 <- glmer(Mittelwert_ng.ml ~ Day_Actual + CI + Weight +
# Season * Beach + (Day_Actual|ID), family = Gamma(link = 'log'), data =
# cortisol.mum.baseline.rsc) --> ## random effects structure is too complex to be
# supported by data

anova(mm1.log, mm1.noID)
```

```
## Data: cortisol.mum.baseline.rsc
## Models:
## mm1.noID: Mittelwert_ng.ml ~ Day_Actual + CI + Weight + Season * Beach
## mm1.log: Mittelwert_ng.ml ~ Day_Actual + CI + Weight + Season * Beach + (1 | ID)
##          npar    AIC    BIC  logLik deviance  Chisq Df Pr(>Chisq)
## mm1.noID    8 723.68 747.49 -353.84   707.68
## mm1.log     9 689.08 715.87 -335.54   671.08 36.604  1  1.447e-09 ***
## ---
## Signif. codes:  0 '***' 0.001 '**' 0.01 '*' 0.05 '.' 0.1 ' ' 1
```

##### Residual check of full model

The residuals of the full model were visually inspected for linearity and equality of error variances (using plots of residuals versus fits), normality (using Q–Q plots) and homogeneity of variance (using Levene's Test).

```r
# check for singularity = some of the constrained parameters of the random
# effects theta parameters are on the boundary
isSingular(mm1.log, tol = 1e-04)
```

```
## [1] FALSE
```

```r
# Check for independence & homoscedasticity - points should be randomly scattered
# around zero for the entire range of fitted values
plot(fitted(mm1.log), resid(mm1.log, type = "pearson"), main = "Residuals vs Fitted")
abline(0, 0, col = "red")
```

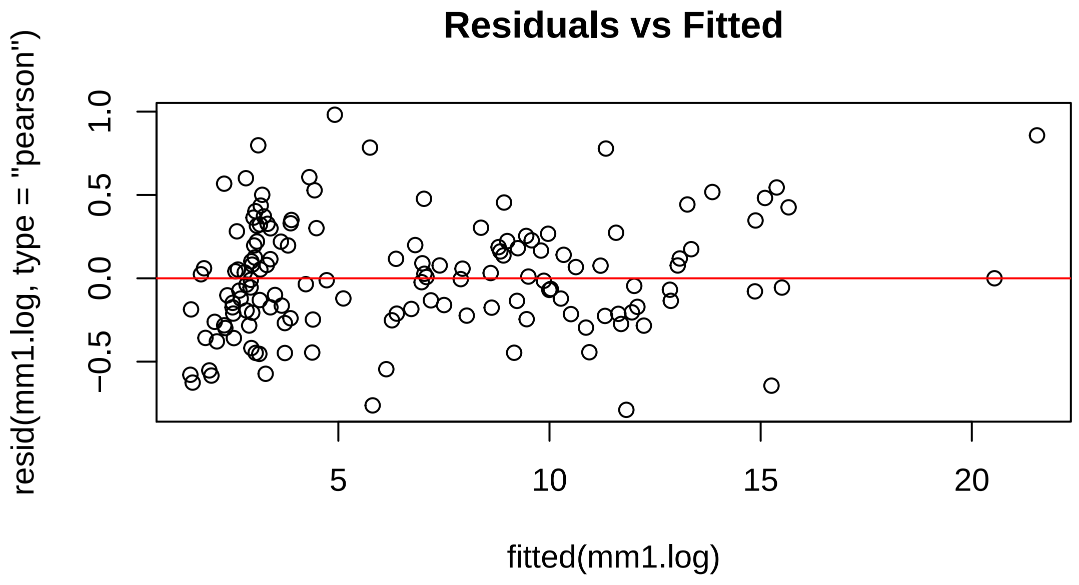

```
# Check if residuals are normally distributed - points should follow the straight
# line.
qqnorm(resid(mm1.log), main = "Normal Q-Q")
qqline(resid(mm1.log), col = "red")
```

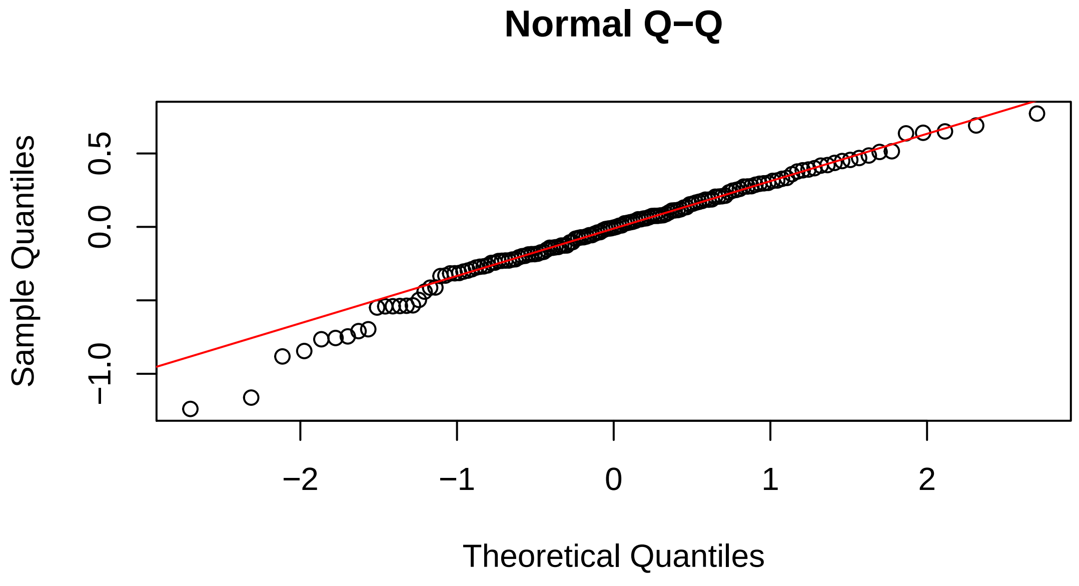

```
# Additional check for homogeneity of variance - Levene's Test 1. Extract the
# residuals and place them in a new column in original data table
cortisol.mum.baseline.rsc$mm1.log.Res <- residuals(mm1.log)
# 2. Create a new column with the absolute value of the residuals
```

```r
cortisol.mum.baseline.rsc$Abs.mm1.log.Res <- abs(cortisol.mum.baseline.rsc$mm1.log.Res)
# 3. Square the absolute values of the residuals to provide more robust estimate
cortisol.mum.baseline.rsc$mm1.log.Res2 <- cortisol.mum.baseline.rsc$Abs.mm1.log.Res^2
# 4. ANOVA of the squared residuals - A p > 0.05 means that the variance of the
# residuals is equal and therefore the assumption of homoscedasticity is met
Levene.Model.F <- lm(mm1.log.Res2 ~ ID, data = cortisol.mum.baseline.rsc)
anova(Levene.Model.F)
```

```
## Analysis of Variance Table
##
## Response: mm1.log.Res2
##           Df Sum Sq  Mean Sq F value Pr(>F)
## ID        91 3.8017 0.041777  0.6975 0.9345
## Residuals 53 3.1744 0.059894
```

##### Model reduction

A backward elimination based on the chi-squared statistic was implemented to simplify the model. The best fit model was determined based on AIC values. The statistical significance of fixed predictors was assessed using a Wald test.

```r
Anova(mm1.log)
```

```
## Analysis of Deviance Table (Type II Wald chisquare tests)
##
## Response: Mittelwert_ng.ml
##                 Chisq Df Pr(>Chisq)
## Day_Actual   267.2607  1  < 2.2e-16 ***
## CI             0.2743  1   0.600478
## Weight         0.5365  1   0.463867
## Season         8.6087  1   0.003346 **
## Beach          3.2481  1   0.071505 .
## Season:Beach   0.0206  1   0.885760
## ---
## Signif. codes:  0 '***' 0.001 '**' 0.01 '*' 0.05 '.' 0.1 ' ' 1
```

```r
mm3 <- update(mm1.log, . ~ . - Season:Beach)
Anova(mm3)
```

```
## Analysis of Deviance Table (Type II Wald chisquare tests)
##
## Response: Mittelwert_ng.ml
##               Chisq Df Pr(>Chisq)
## Day_Actual 269.0156  1  < 2.2e-16 ***
## CI           0.2659  1   0.606125
## Weight       0.5300  1   0.466589
## Season       8.6068  1   0.003349 **
## Beach        3.2471  1   0.071552 .
## ---
## Signif. codes:  0 '***' 0.001 '**' 0.01 '*' 0.05 '.' 0.1 ' ' 1
```

```
mm4 <- update(mm3, . ~ . - CI)
Anova(mm4)
```

```
## Analysis of Deviance Table (Type II Wald chisquare tests)
##
## Response: Mittelwert_ng.ml
##                 Chisq Df Pr(>Chisq)
## Day_Actual 341.1368  1  < 2.2e-16 ***
## Weight       0.3563  1   0.550547
## Season       8.4679  1   0.003615 **
## Beach        3.3514  1   0.067147 .
## ---
## Signif. codes:  0 '***' 0.001 '**' 0.01 '*' 0.05 '.' 0.1 ' ' 1
```

```
mm5 <- update(mm4, . ~ . - Weight)
Anova(mm5)
```

```
## Analysis of Deviance Table (Type II Wald chisquare tests)
##
## Response: Mittelwert_ng.ml
##                 Chisq Df Pr(>Chisq)
## Day_Actual 347.0147  1  < 2.2e-16 ***
## Season       8.2469  1   0.004082 **
## Beach        3.3257  1   0.068204 .
## ---
## Signif. codes:  0 '***' 0.001 '**' 0.01 '*' 0.05 '.' 0.1 ' ' 1
```

```
mm6 <- update(mm5, . ~ . - Beach)
Anova(mm6)
```

```
## Analysis of Deviance Table (Type II Wald chisquare tests)
##
## Response: Mittelwert_ng.ml
##                 Chisq Df Pr(>Chisq)
## Day_Actual 338.9563  1  < 2.2e-16 ***
## Season       8.0214  1   0.004623 **
## ---
## Signif. codes:  0 '***' 0.001 '**' 0.01 '*' 0.05 '.' 0.1 ' ' 1
```

```
mm7 <- update(mm6, . ~ . - Season)
Anova(mm7)
```

```
## Analysis of Deviance Table (Type II Wald chisquare tests)
##
## Response: Mittelwert_ng.ml
##               Chisq Df Pr(>Chisq)
## Day_Actual 323.88  1  < 2.2e-16 ***
## ---
## Signif. codes:  0 '***' 0.001 '**' 0.01 '*' 0.05 '.' 0.1 ' ' 1
```

```r
anova(mm1.log, mm4, mm5, mm6, mm7)
```

```
## Data: cortisol.mum.baseline.rsc
## Models:
## mm7: Mittelwert_ng.ml ~ Day_Actual + (1 | ID)
## mm6: Mittelwert_ng.ml ~ Day_Actual + Season + (1 | ID)
## mm5: Mittelwert_ng.ml ~ Day_Actual + Season + Beach + (1 | ID)
## mm4: Mittelwert_ng.ml ~ Day_Actual + Weight + Season + Beach + (1 | ID)
## mm1.log: Mittelwert_ng.ml ~ Day_Actual + CI + Weight + Season * Beach + (1 | ID)
##         npar    AIC    BIC  logLik deviance  Chisq Df Pr(>Chisq)
## mm7        4 690.64 702.54 -341.32   682.64
## mm6        5 684.98 699.86 -337.49   674.98 7.6597  1   0.005647 **
## mm5        6 683.72 701.58 -335.86   671.72 3.2590  1   0.071034 .
## mm4        7 685.36 706.20 -335.68   671.36 0.3535  1   0.552130
## mm1.log    9 689.08 715.87 -335.54   671.08 0.2867  2   0.866455
## ---
## Signif. codes:  0 '***' 0.001 '**' 0.01 '*' 0.05 '.' 0.1 ' ' 1
```

```r
summary(mm5)
```

```
## Generalized linear mixed model fit by maximum likelihood (Laplace
##   Approximation) [glmerMod]
##  Family: Gamma  ( log )
## Formula: Mittelwert_ng.ml ~ Day_Actual + Season + Beach + (1 | ID)
##    Data: cortisol.mum.baseline.rsc
##
##      AIC      BIC   logLik deviance df.resid
##    683.7    701.6   -335.9    671.7      139
##
## Scaled residuals:
##      Min       1Q   Median       3Q      Max
## -1.96602 -0.52211 -0.01456  0.58909  2.53562
##
## Random effects:
##  Groups   Name        Variance Std.Dev.
##  ID       (Intercept) 0.1230   0.3508
##  Residual             0.1622   0.4027
## Number of obs: 145, groups:  ID, 92
##
## Fixed effects:
##              Estimate Std. Error t value Pr(>|z|)
## (Intercept)  1.38843    0.10737  12.932  < 2e-16 ***
## Day_Actual  -0.65990    0.03542 -18.628  < 2e-16 ***
## Season1920   0.35353    0.12310   2.872  0.00408 **
## BeachSSB     0.22434    0.12302   1.824  0.06820 .
## ---
## Signif. codes:  0 '***' 0.001 '**' 0.01 '*' 0.05 '.' 0.1 ' ' 1
##
## Correlation of Fixed Effects:
##            (Intr) Dy_Act Ss1920
## Day_Actual  0.125
## Season1920 -0.573 -0.106
## BeachSSB   -0.584 -0.103 -0.009
```

```r
# Check model for Multicollinearity
performance::check_collinearity(mm5)
```

```
## # Check for Multicollinearity
##
## Low Correlation
##
##    Parameter  VIF Increased SE
##   Day_Actual 1.02         1.01
##       Season 1.01         1.01
##        Beach 1.01         1.01
```

```r
# variance estimates: variance of ID (Intercept) = variability of the intercept
# across ID & variance of ID (day) = variability in the slope across ID &
# residual variance = s2 = residual variation around the regression line
vc <- lme4::VarCorr(mm5)
print(vc, comp = c("Variance", "Std.Dev."), digits = 2)
```

```
##  Groups   Name        Variance Std.Dev.
##  ID       (Intercept) 0.12     0.35
##  Residual             0.16     0.40
```

```r
# Marginal and conditional R2
r.squaredGLMM(mm5)
```

```
##                 R2m       R2c
## delta     0.6033827 0.7744911
## lognormal 0.6135187 0.7875014
## trigamma  0.5919764 0.7598501
```

```r
# ICC = % of the variation in outcome attributable to differences between random
# effect (subjects)
icc_specs(mm5, percent = TRUE)
```

```
##        grp      vcov       icc  percent
## 1       ID 0.1230459 0.4314193 43.14193
## 2 Residual 0.1621660 0.5685807 56.85807
```

### Animal models

#### Pedigree animal model

To estimate the amount of baseline cortisol variance that is due to genetic differences among individuals, we built two animal models. The first included a simple pedigree to estimate the additive source of genetic variance. The pedigree was included as a random effect. This allows us to estimate the amount of variance explained by genetics, a source of non-independence in the population. The posterior distribution of the model intercept and autocorrelation was checked to assess model fit.

```
#### build pedigree ----
ped <- cortisol[!duplicated(cortisol$ID), ]
ped <- subset(ped, select = c("ID", "ID_status", "ID_pair"))
ped <- ped[order(ped$ID_status), ]
ped$FATHER <- NA
ped$MOTHER <- ifelse(ped$ID_status == "mum", NA, ped$ID_pair)
ped <- subset(ped, select = -c(ID_status, ID_pair))

#### prepare data for animal model ----
ped$id <- as.factor(ped$ID)
ped$father <- as.factor(ped$FATHER)
ped$mother <- as.factor(ped$MOTHER)
ped <- subset(ped, select = -c(ID, MOTHER, FATHER))
ped <- GeneticsPed::extend(ped)
ped <- as.data.frame(ped)

cortisol.animalmodel <- cortisol
names(cortisol.animalmodel)[names(cortisol.animalmodel) == "ID"] <- "animal"
cortisol.animalmodel$animal <- as.factor(cortisol.animalmodel$animal)

cortisol.animalmodel$MOTHER <- ifelse(cortisol.animalmodel$ID_status == "mum", NA,
    cortisol.animalmodel$ID_pair)
cortisol.animalmodel$MOTHER <- as.factor(cortisol.animalmodel$MOTHER)

#### animal model with simple pedigree ----
cortisol.animalmodel.baseline <-
    cortisol.animalmodel[cortisol.animalmodel$SampleCollected == "Baseline", ]

p.var <- var(cortisol.animalmodel.baseline$Mittelwert_ng.ml, na.rm = TRUE)
prior1.2 <- list(G = list(G1 = list(V = matrix(p.var * 0.05), nu = 1)), R = list(V =
    matrix(p.var * 0.95), nu = 1))

m.animal.2 <- MCMCglmm(Mittelwert_ng.ml ~ 1, random = ~animal, pedigree = ped, data =
    cortisol.animalmodel.baseline, prior = prior1.2, nitt = 9e+06, thin = 8500, burnin =
    15000, verbose = FALSE)

# shows distributions of the estimates of the additive genetic (animal) and
# residual (units) effects - should see a consistent amount of variation around a
# largely unchanging mean value of the intercept
plot(m.animal.2$VCV)
```

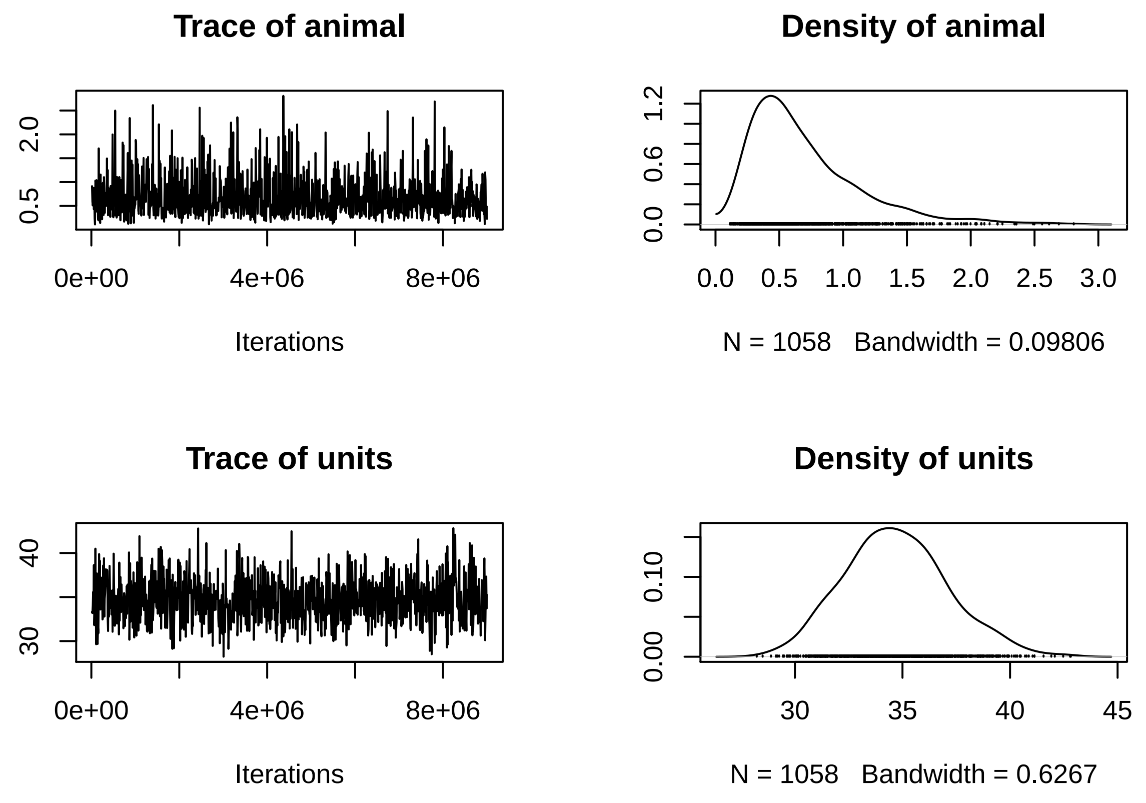

```r
summary(m.animal.2)
```

```
##
##  Iterations = 15001:8999501
##  Thinning interval  = 8500
##  Sample size  = 1058
##
##  DIC: 2805.462
##
##  G-structure:  ~animal
##
##        post.mean l-95% CI u-95% CI eff.samp
## animal    0.6976   0.1122    1.551     1058
##
##  R-structure:  ~units
##
##       post.mean l-95% CI u-95% CI eff.samp
## units      34.7    30.61    39.95     1058
##
##  Location effects: Mittelwert_ng.ml ~ 1
##
##             post.mean l-95% CI u-95% CI eff.samp  pMCMC
## (Intercept)     7.714    7.099    8.261     1058 <9e-04 ***
## ---
## Signif. codes:  0 '***' 0.001 '**' 0.01 '*' 0.05 '.' 0.1 ' ' 1
```

```r
# estimates of the additive genetic and residual variance by calculating the
# modes of the posterior distributions = an estimate of the proportion of
# variance in cortisol explained by additive effects
posterior.mode(m.animal.2$VCV)
```

```
##     animal      units
##  0.4954163 34.6633896
```

```r
# Bayesian equivalent of confidence intervals by calculating the values of the
# estimates that bound 95% of the posterior distributions:
HPDinterval(m.animal.2$VCV)
```

```
##             lower     upper
## animal  0.1121572  1.551081
## units  30.6140690 39.949569
## attr(,"Probability")
## [1] 0.9499055
```

```r
# estimating heritability
posterior.heritability <- m.animal.2$VCV[, "animal"]/(m.animal.2$VCV[, "animal"] +
    m.animal.2$VCV[, "units"])
HPDinterval(posterior.heritability, 0.95) * 100 # in percent
```

```
##          lower    upper
## var1 0.4096818 4.476451
## attr(,"Probability")
## [1] 0.9499055
```

```r
posterior.mode(posterior.heritability) * 100 # in percent
```

```
##     var1
## 1.330326
```

```r
plot(posterior.heritability)
```

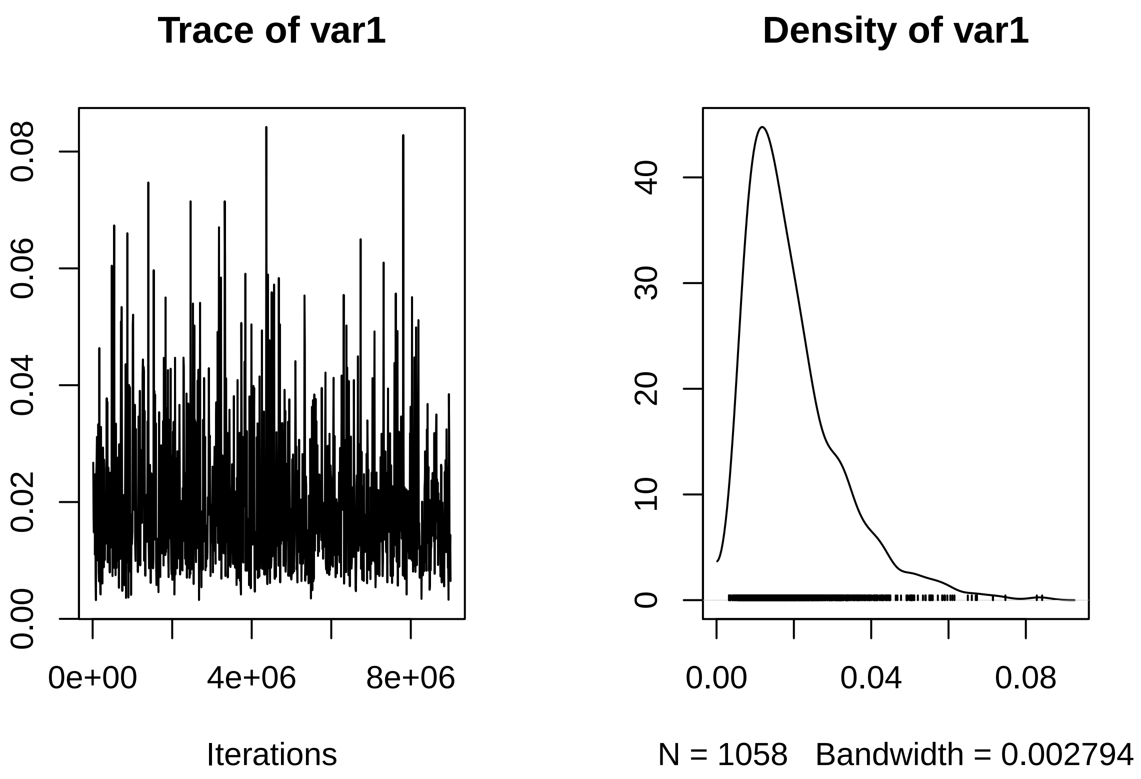

##### SNP relatedness matrix animal model

The second animal model included a relatedness matrix built from 72,354 SNPs for the n = 98 individuals sampled in the 2018-2019 season. The relatedness matrix was included as a random effect. The posterior distribution of the model intercept and autocorrelation was checked to assess model fit.

```
#### prepare data for animal model ----

# The identities of individuals in the phenotypic data set (cortisol) are linked
# to the inverse of the relatedness matrix by including it in the 'ginverse', and
# the random effect is specified by 'dom', which is identical to the animal
# column
cortisol.animalmodel$dom <- cortisol.animalmodel$animal

##### filter dataset

# no SNP data for second season - remove from cortisol dataframe
cortisol.animalmodel.snp <- cortisol.animalmodel[cortisol.animalmodel$Season == "1819", ]

# no genetic data for F9 - remove from cortisol dataframe
cortisol.animalmodel.snp <- cortisol.animalmodel.snp[!cortisol.animalmodel.snp$dom ==
    "F9", ]

# no cortisol data for C4, F26, F4 - remove from matrix
row.names.remove <- c("F26", "F4")
```

```
snp.matrix <- snp.matrix[!(row.names(snp.matrix) %in% row.names.remove), ]
snp.matrix <- subset(snp.matrix, select = -c(F26, F4))

# convert relatedness matrix in dgCMatrix format
snp.dgCmatrix <- as(solve(snp.matrix), "dgCMatrix")

#### animal model with SNP relatedness matrix ----
prior1.3.1 <- list(G = list(G1 = list(V = matrix(p.var * 0.05), nu = 1), G2 = list(V =
    matrix(p.var * 0.05), nu = 1)), R = list(V = matrix(p.var * 0.95), nu = 1))

m.animal.3.1 <- MCMCglmm(Mittelwert_ng.ml ~ 1, random = ~animal + dom, ginverse =
    list(dom = snp.dgCmatrix), data = cortisol.animalmodel.snp, prior = prior1.3.1, nitt
    = 9e+06, thin = 8500, burnin = 15000, verbose = FALSE)

plot(m.animal.3.1$VCV)
```

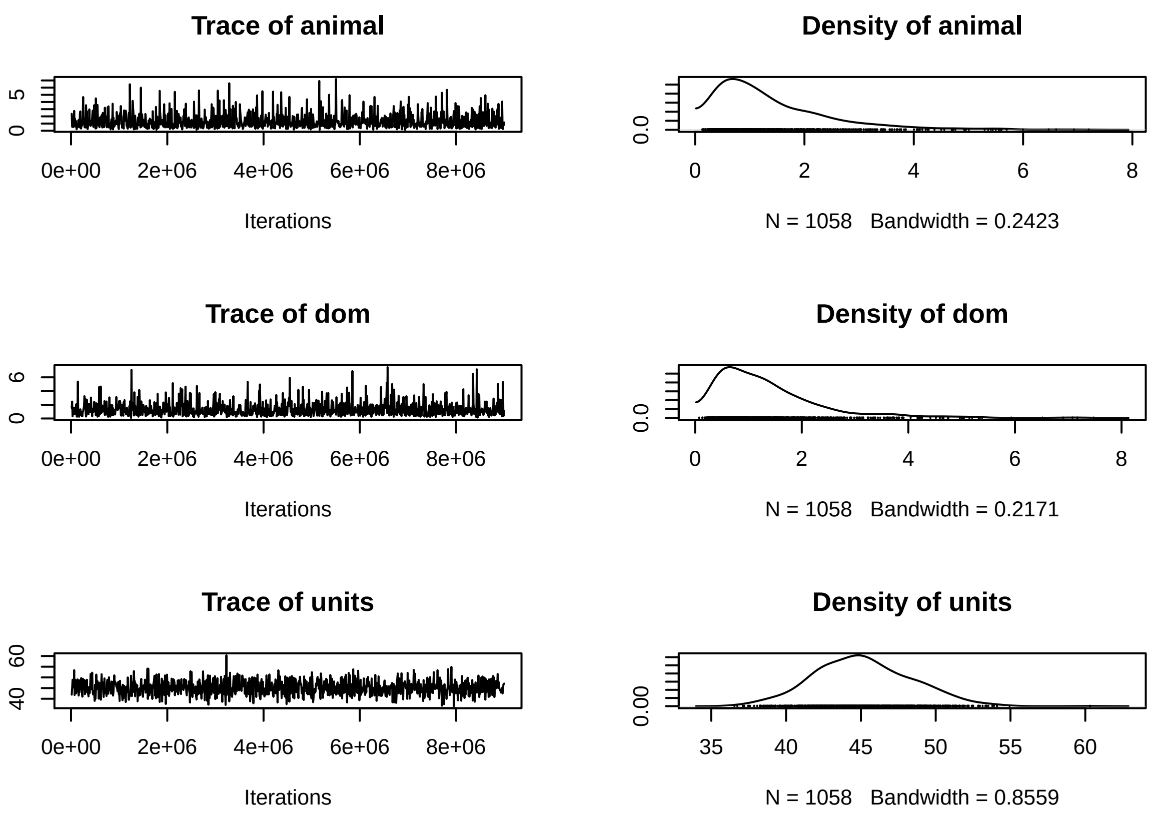

```
posterior.mode(m.animal.3.1$VCV)
```

```
##     animal        dom      units
##  0.9135748  0.5049596 44.5704907
```

```
HPDinterval(m.animal.3.1$VCV)
```

```
##              lower     upper
```

```
## animal  0.1386852  3.563790
## dom     0.1913167  3.613973
## units  38.9930790 51.626551
## attr(,"Probability")
## [1] 0.9499055
```

```
# Calculate posterior heritability and the associated 95% confidence intervals
posterior.heritability.3.1 <- m.animal.3.1$VCV[, "animal"]/(m.animal.3.1$VCV[, "animal"]
    + m.animal.3.1$VCV[, "units"] + m.animal.3.1$VCV[, "dom"])

HPDinterval(posterior.heritability.3.1, 0.95) * 100 # in percent
```

```
##          lower    upper
## var1 0.2997903 7.218844
## attr(,"Probability")
## [1] 0.9499055
```

```
posterior.mode(posterior.heritability.3.1) * 100 # in percent
```

```
##     var1
## 1.497993
```

```
plot(posterior.heritability.3.1)
```

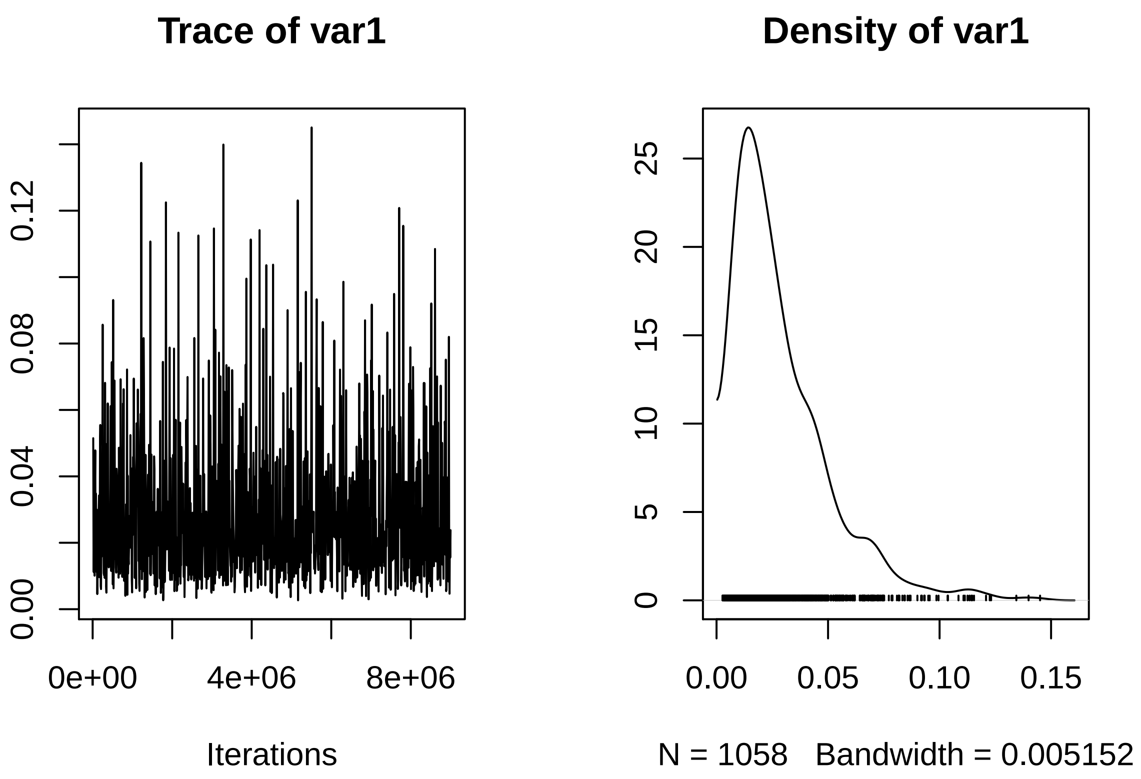

#### Manuscript figures

```
m3.CI <- data.frame(stringsAsFactors = FALSE, Predictors = c("(Intercept)", "Age", "Body
    condition", "Weight", "Season [2020]", "Colony [SSB]", "Sex [male]"), Estimates =
    c(1.58, -0.29, 0.14, -0.36, 0.25, 0.13, 0.24), CIupper = c(1.75, -0.14, 0.31, -0.2,
    0.41, 0.29, 0.41), CIlower = c(1.4, -0.44, -0.02, -0.52, 0.09, -0.03, 0.08))

m3.CI$Predictors <- factor(m3.CI$Predictors, levels = c("Colony [SSB]", "Body condition",
    "Season [2020]", "Sex [male]", "Weight", "Age", "(Intercept)"))

ggplot(data = m3.CI, aes(x = Estimates, y = Predictors)) + geom_point() +
    geom_errorbar(aes(xmin = CIlower, xmax = CIupper), width = 0.1) +
    geom_vline(xintercept = 0, linetype = "solid", color = "black", size = 0.5) +
    theme_classic() + theme(text = element_text(size = 15), legend.position = "none",
    plot.margin = unit(c(10, 5, 5, 0), "mm"), plot.title = element_text(margin = margin(b
    = 0), size = 15), plot.subtitle = element_text(hjust = 0.28, margin = margin(t = 5, b
    = 10), size = 12)) + labs(x = "Estimates", y = "", title = "Pup baseline cortisol",
    subtitle = "") + scale_x_continuous(minor_breaks = NULL, limits = c(-0.8, 1.75),
    breaks = c(-0.5, 0, 0.5, 1, 1.5), labels = c(-0.5, 0, 0.5, 1, 1.5))
```

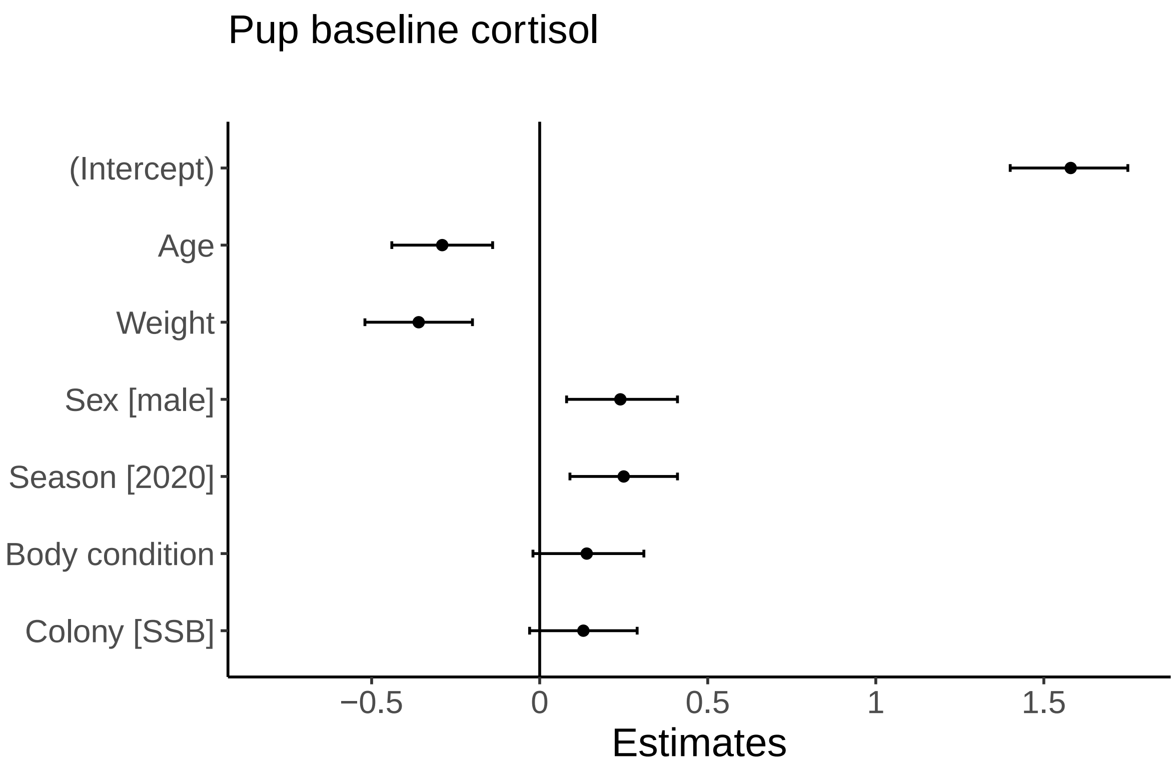

```
ggplot() + geom_point(data = cortisol.pup.baseline, aes(y = Mittelwert_ng.ml, x =
    Weight)) + geom_smooth(data = cortisol.pup.baseline[cortisol.pup.baseline$Day == 0,
    ], aes(y = Mittelwert_ng.ml, x = Weight, colour = "#d95f02"), formula = y ~ x, method
    = "lm", se = FALSE, size = 2, fullrange = F) + geom_smooth(data =
    cortisol.pup.baseline[cortisol.pup.baseline$Day == 20, ], aes(y = Mittelwert_ng.ml, x
    = Weight, colour = "#e7298a"), formula = y ~ x, method = "lm", se = FALSE, size = 2,
    fullrange = F) + geom_smooth(data = cortisol.pup.baseline[cortisol.pup.baseline$Day
    == 40, ], aes(y = Mittelwert_ng.ml, x = Weight, colour = "#66a61e"), formula = y ~ x,
    method = "lm", se = FALSE, size = 2, fullrange = F) + geom_smooth(data =
    cortisol.pup.baseline[cortisol.pup.baseline$Day >= 41, ], aes(y = Mittelwert_ng.ml, x
    = Weight, colour = "#e6ab02"), formula = y ~ x, method = "lm", se = FALSE, size = 2,
    fullrange = F) + scale_color_identity(guide = "legend", name = "Slope", breaks =
    c("#d95f02", "#e7298a", "#66a61e", "#e6ab02"), labels = c("birth", "20 days old", "40
    days old", "molt")) + theme_classic() + theme(text = element_text(size = 15),
    plot.margin = unit(c(10, 5, 5, 5), "mm"), legend.key.size = unit(2, "mm"),
    legend.title = element_text(size = 12), legend.position = c(0.8, 0.8), plot.title =
    element_text(margin = margin(b = 0), size = 15), plot.subtitle = element_text(hjust =
    0.28, margin = margin(t = 5, b = 10), size = 12)) + labs(x = "Weight (kg)", y =
    "Baseline (ng/mL)", title = "Pup baseline cortisol", subtitle = "")
```

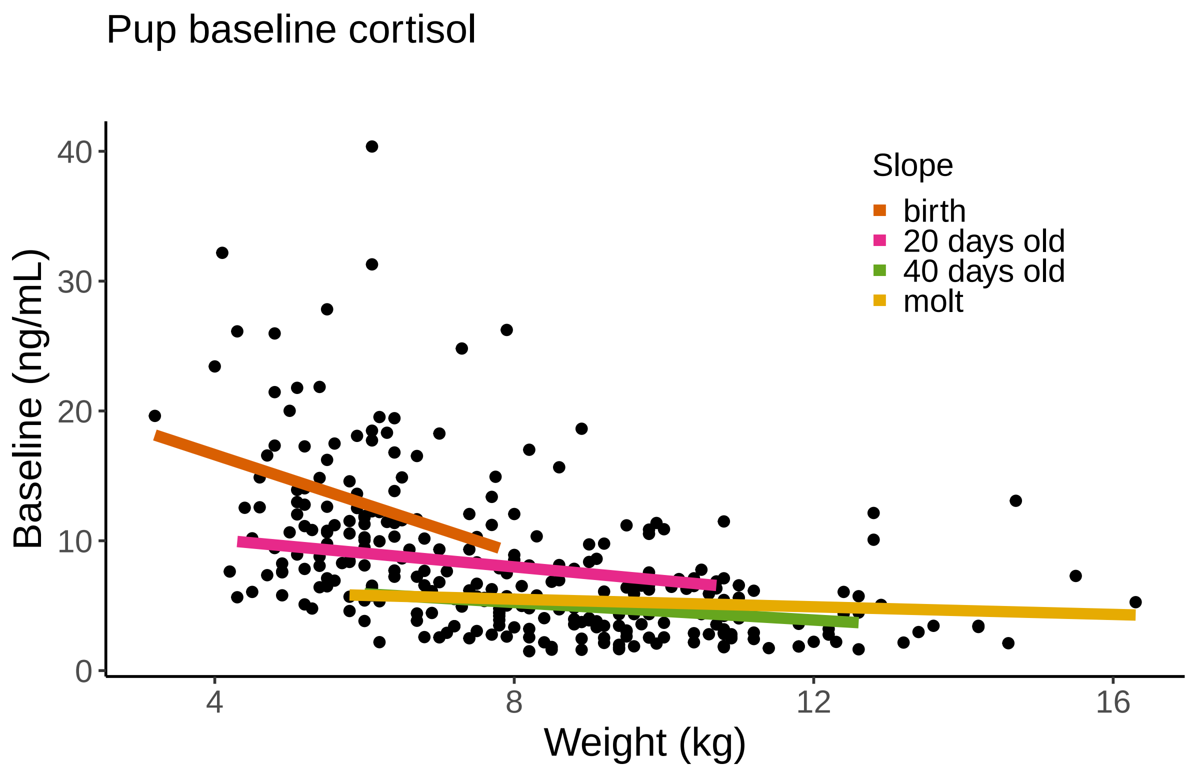

```
ggplot(data = cortisol.pup.baseline) + geom_boxplot(aes(y = Mittelwert_ng.ml, x = Day,
    group = Day), fill = "#ffffff", na.rm = TRUE, outlier.shape = NA) + geom_point(aes(y
    = Mittelwert_ng.ml, x = Day_Actual, group = Day), na.rm = TRUE, alpha = 0.25) +
    theme_classic() + theme(text = element_text(size = 15), legend.position = "none",
    plot.margin = unit(c(10, 5, 5, 5), "mm"), plot.title = element_text(margin = margin(b
    = 0), size = 15), plot.subtitle = element_text(hjust = 0.28, margin = margin(t = 5, b
    = 10), size = 12)) + labs(x = "Age (days)", y = "Baseline (ng/mL)", title = "Pup
    baseline cortisol", subtitle = "") + scale_x_continuous(minor_breaks = NULL, breaks =
    c(0, 20, 40, 60, 80), labels = c(0, 20, 40, 60, 80))
```

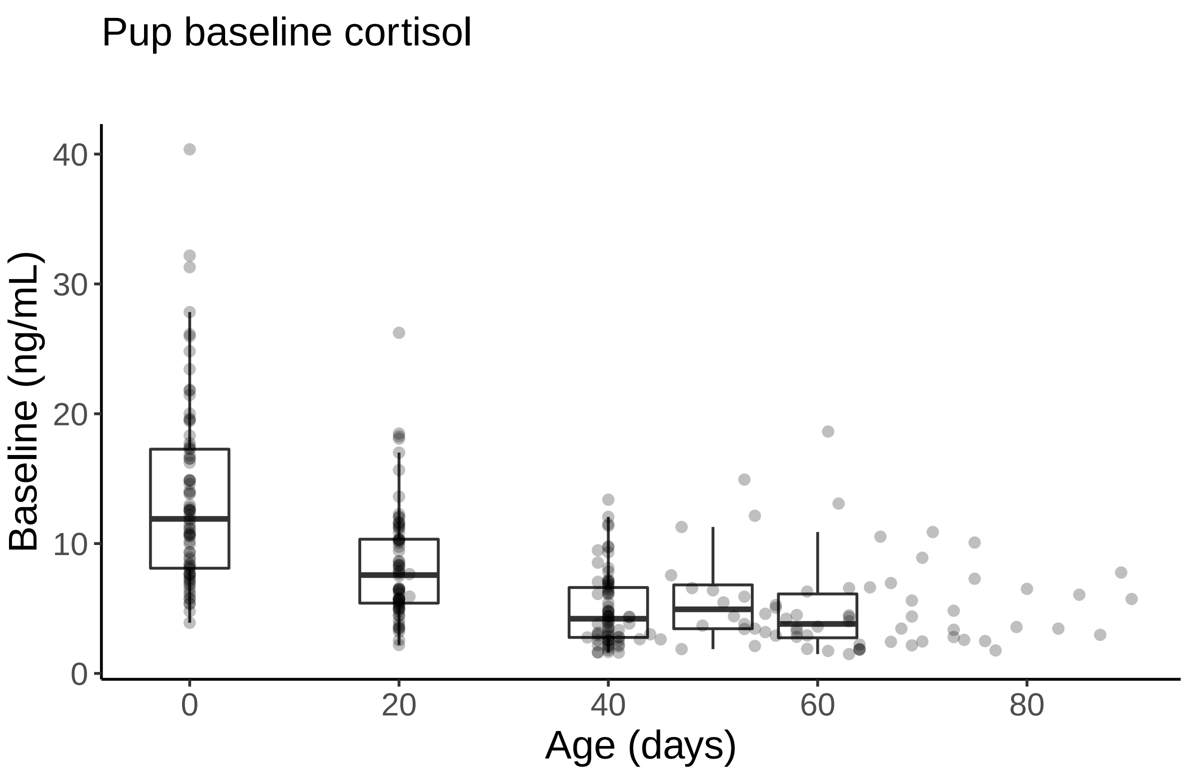

```
ggplot(data = cortisol.pup.baseline) + geom_boxplot(aes(y = Mittelwert_ng.ml, x = Season,
    group = Season), na.rm = TRUE, outlier.shape = NA, fill = c("#bcbcbc", "#515151")) +
    geom_point(aes(y = Mittelwert_ng.ml, x = Season), na.rm = TRUE, alpha = 0.25) +
    theme_classic() + theme(text = element_text(size = 15), legend.position = "none",
    plot.margin = unit(c(10, 5, 5, 5), "mm"), plot.title = element_text(margin = margin(b
    = 0), size = 15), plot.subtitle = element_text(hjust = 0.28, margin = margin(t = 5, b
    = 10), size = 12)) + labs(x = "Year", y = "Baseline (ng/mL)", title = "Pup baseline
    cortisol", subtitle = "") + scale_x_discrete(breaks = c("1819", "1920"), labels =
    c("2019", "2020"))
```

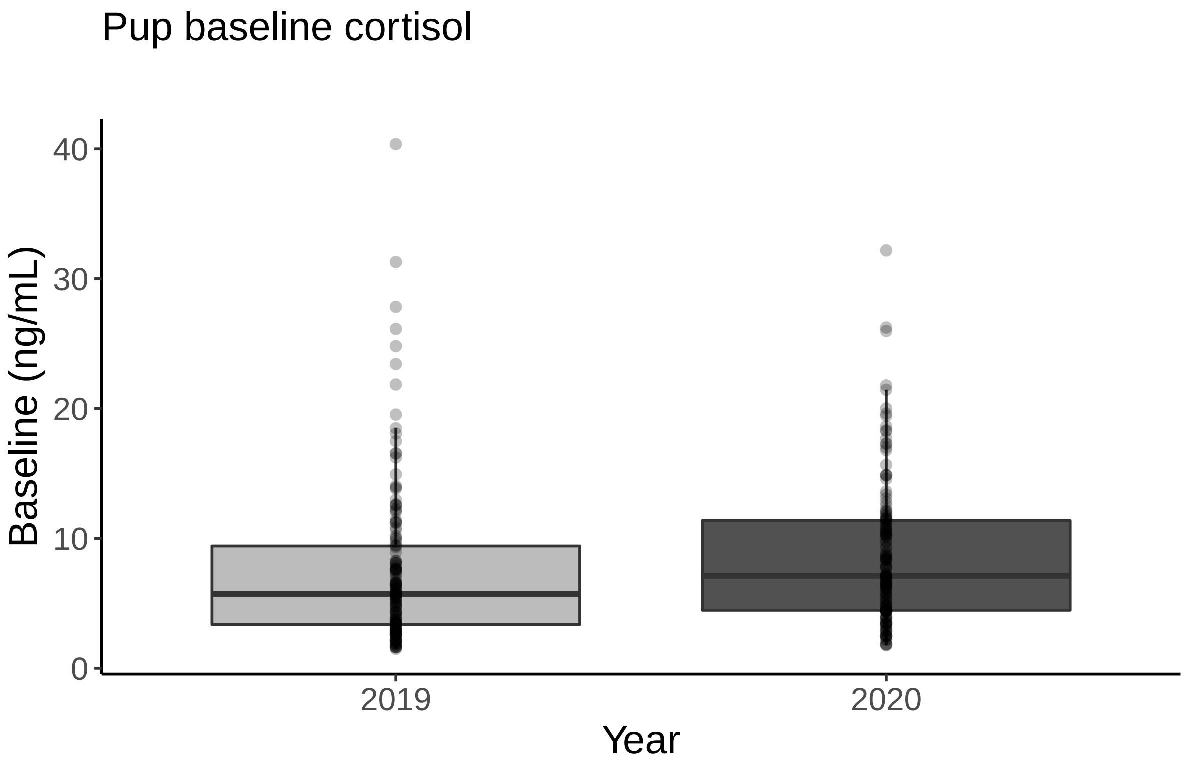

```r
ggplot(data = cortisol.pup.baseline) + geom_boxplot(aes(y = Mittelwert_ng.ml, x = Sex,
    group = Sex), na.rm = TRUE, outlier.shape = NA, fill = "#ffffff") + geom_point(aes(y
    = Mittelwert_ng.ml, x = Sex), na.rm = TRUE, alpha = 0.25) + theme_classic() +
    theme(text = element_text(size = 15), legend.position = "none", plot.margin =
    unit(c(10, 5, 5, 5), "mm"), plot.title = element_text(margin = margin(b = 0), size =
    15), plot.subtitle = element_text(hjust = 0.28, margin = margin(t = 5, b = 10), size
    = 12)) + labs(x = "Sex", y = "Baseline (ng/mL)", title = "Pup baseline cortisol",
    subtitle = "") + scale_x_discrete(breaks = c("F", "M"), labels = c("Female", "Male"))
```

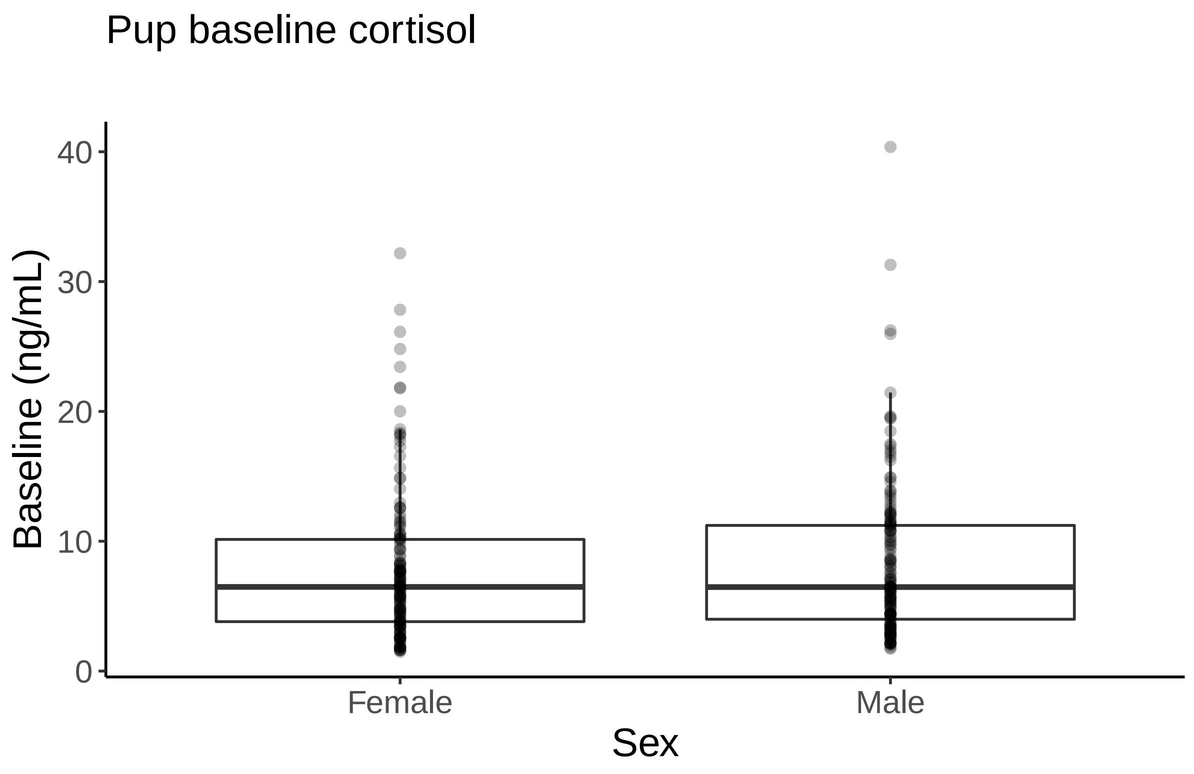

```
mm5.CI <- data.frame(stringsAsFactors = FALSE, Predictors = c("(Intercept)", "Days
    postpartum", "Season [2020]", "Colony [SSB]"), Estimates = c(1.39, -0.66, 0.35,
    0.22), CIupper = c(1.6, -0.59, 0.59, 0.47), CIlower = c(1.18, -0.73, 0.11, -0.02))

mm5.CI$Predictors <- factor(mm5.CI$Predictors, levels = c("Colony [SSB]", "Season
    [2020]", "Days postpartum", "(Intercept)"))

ggplot(data = mm5.CI, aes(x = Estimates, y = Predictors)) + geom_point() +
    geom_errorbar(aes(xmin = CIlower, xmax = CIupper), width = 0.1) +
    geom_vline(xintercept = 0, linetype = "solid", color = "black", size = 0.5) +
    theme_classic() + theme(text = element_text(size = 15), legend.position = "none",
    plot.margin = unit(c(10, 5, 5, 0), "mm"), plot.title = element_text(margin = margin(b
    = 0), size = 15), plot.subtitle = element_text(hjust = 0.28, margin = margin(t = 5, b
    = 10), size = 12)) + labs(x = "Estimates", y = "", title = "Maternal baseline
    cortisol", subtitle = "") + scale_x_continuous(minor_breaks = NULL, limits = c(-0.8,
    1.75), breaks = c(-0.5, 0, 0.5, 1, 1.5), labels = c(-0.5, 0, 0.5, 1, 1.5))
```

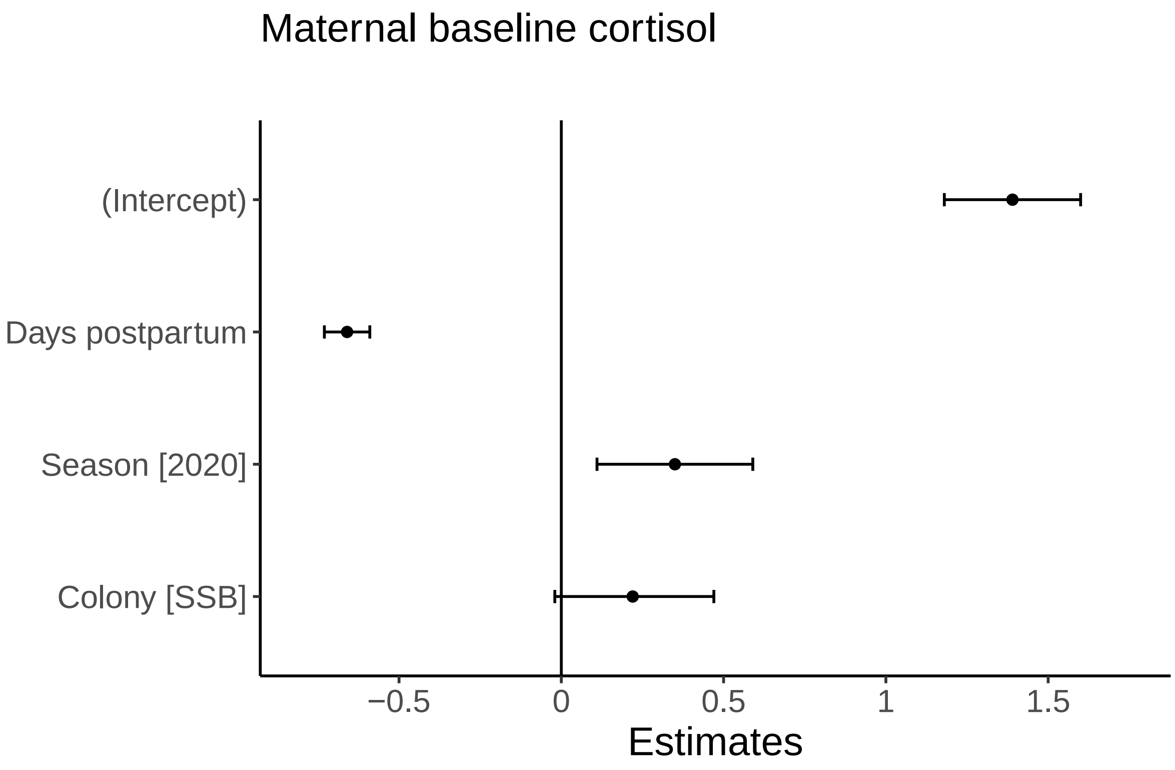

```
ggplot() + geom_boxplot(data = cortisol.mum.baseline[cortisol.mum.baseline$Day == 0, ],
    aes(y = Mittelwert_ng.ml, x = Day), fill = "#ffffff", na.rm = TRUE, outlier.shape =
    NA, width = 10) + geom_point(data = cortisol.mum.baseline, aes(y = Mittelwert_ng.ml,
    x = Day_Actual), na.rm = TRUE, alpha = 0.25) + theme_classic() + theme(text =
    element_text(size = 15), legend.position = "none", plot.margin = unit(c(10, 5, 5, 5),
    "mm"), plot.title = element_text(margin = margin(b = 0), size = 15), plot.subtitle =
    element_text(hjust = 0.28, margin = margin(t = 5, b = 10), size = 12)) + labs(x =
    "Days postpartum", y = "Baseline (ng/mL)", title = "Maternal baseline cortisol",
    subtitle = "") + scale_x_continuous(minor_breaks = NULL, breaks = c(0, 20, 40, 60,
    80), labels = c(0, 20, 40, 60, 80))
```

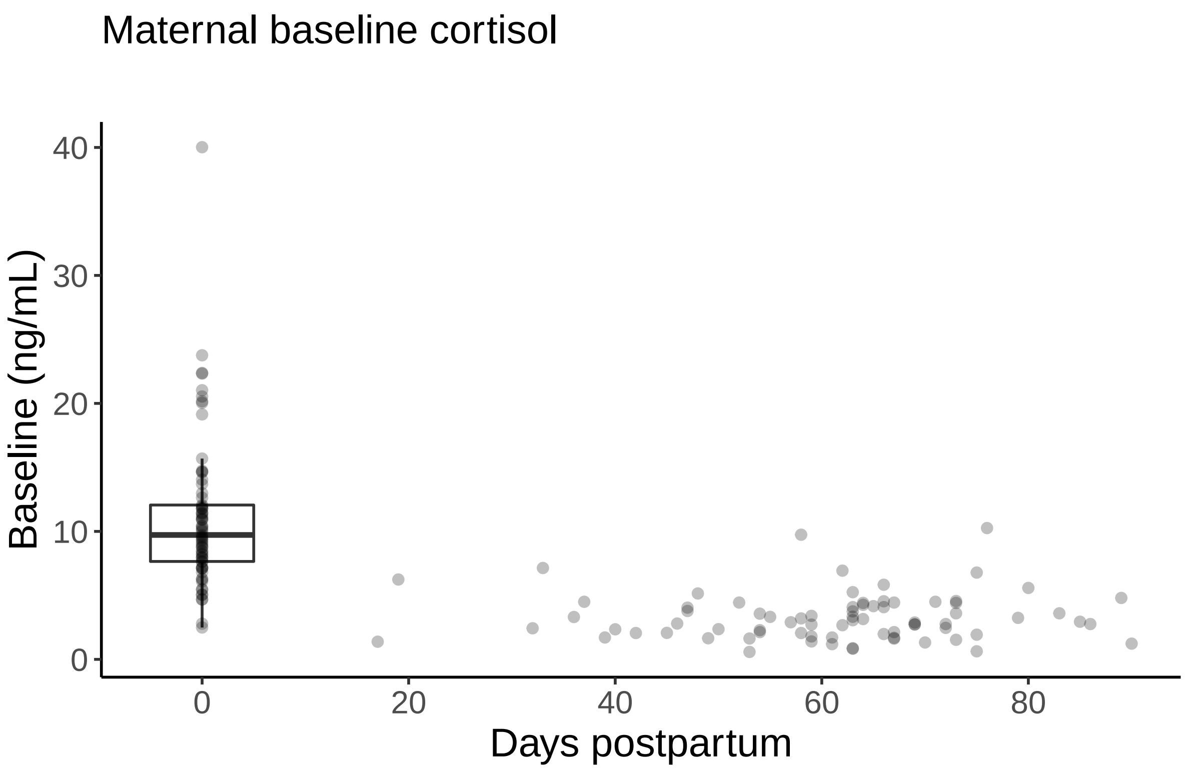

```
ggplot(data = cortisol.mum.baseline) + geom_boxplot(aes(y = Mittelwert_ng.ml, x = Season,
    group = Season), na.rm = TRUE, outlier.shape = NA, fill = c("#bcbcbc", "#515151")) +
    geom_point(aes(y = Mittelwert_ng.ml, x = Season), na.rm = TRUE, alpha = 0.25) +
    theme_classic() + theme(text = element_text(size = 15), legend.position = "none",
    plot.margin = unit(c(10, 5, 5, 5), "mm"), plot.title = element_text(margin = margin(b
    = 0), size = 15), plot.subtitle = element_text(hjust = 0.28, margin = margin(t = 5, b
    = 10), size = 12)) + labs(x = "Year", y = "Baseline (ng/mL)", title = "Maternal
    baseline cortisol", subtitle = "") + scale_x_discrete(breaks = c("1819", "1920"),
    labels = c("2019", "2020"))
```

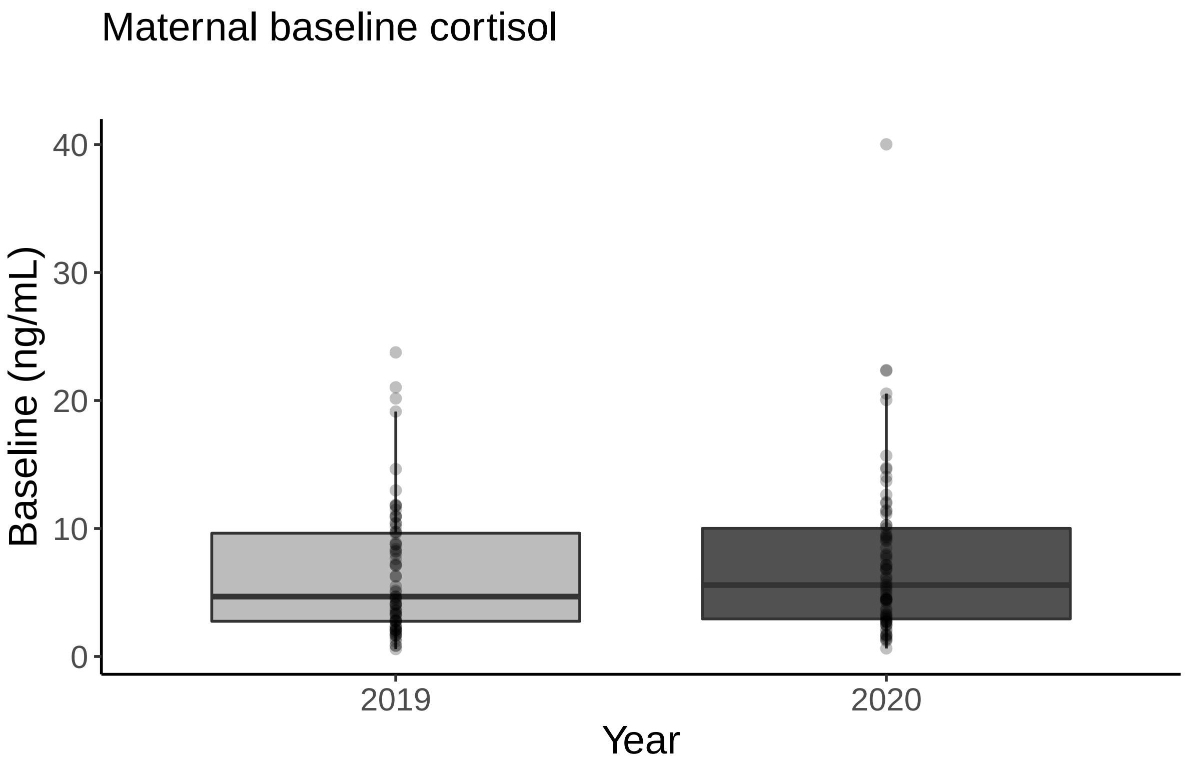

```r
posterior.heritability.pedigree.snps <- data.frame(Pedigree =
    posterior.heritability[1:1058], SNP = posterior.heritability.3.1[1:1058])

posterior.heritability.pedigree.snps.long <-
    melt(setDT(posterior.heritability.pedigree.snps), variable.name = "model", value.name
    = "estimate")

posterior.mode.pedigree.snps <- data.frame(model = c("Pedigree", "SNP"), mode =
    c(posterior.mode(posterior.heritability),
    posterior.mode(posterior.heritability.3.1)), lower =
    c(HPDinterval(posterior.heritability, 0.95)[1],
    HPDinterval(posterior.heritability.3.1, 0.95)[1]), upper =
    c(HPDinterval(posterior.heritability, 0.95)[2],
    HPDinterval(posterior.heritability.3.1, 0.95)[2]))

ggplot() + geom_sina(data = posterior.heritability.pedigree.snps.long, aes(x = model, y =
    estimate), alpha = 0.3, cex = 2) + geom_point(data = posterior.mode.pedigree.snps,
    aes(x = model, y = mode), col = "#E69F00", size = 5) + geom_segment(data =
    posterior.mode.pedigree.snps, aes(x = model, xend = model, y = lower, yend = upper),
    col = "#E69F00", size = 2) + theme_classic() + theme(text = element_text(size = 15),
    plot.title = element_text(size = 15)) + labs(x = " ", y = "Heritability estimate\n",
    title = "(a) Baseline cortisol") + scale_x_discrete(breaks = c("Pedigree", "SNP"),
    labels = c("Pedigree model", "SNP model"))
```

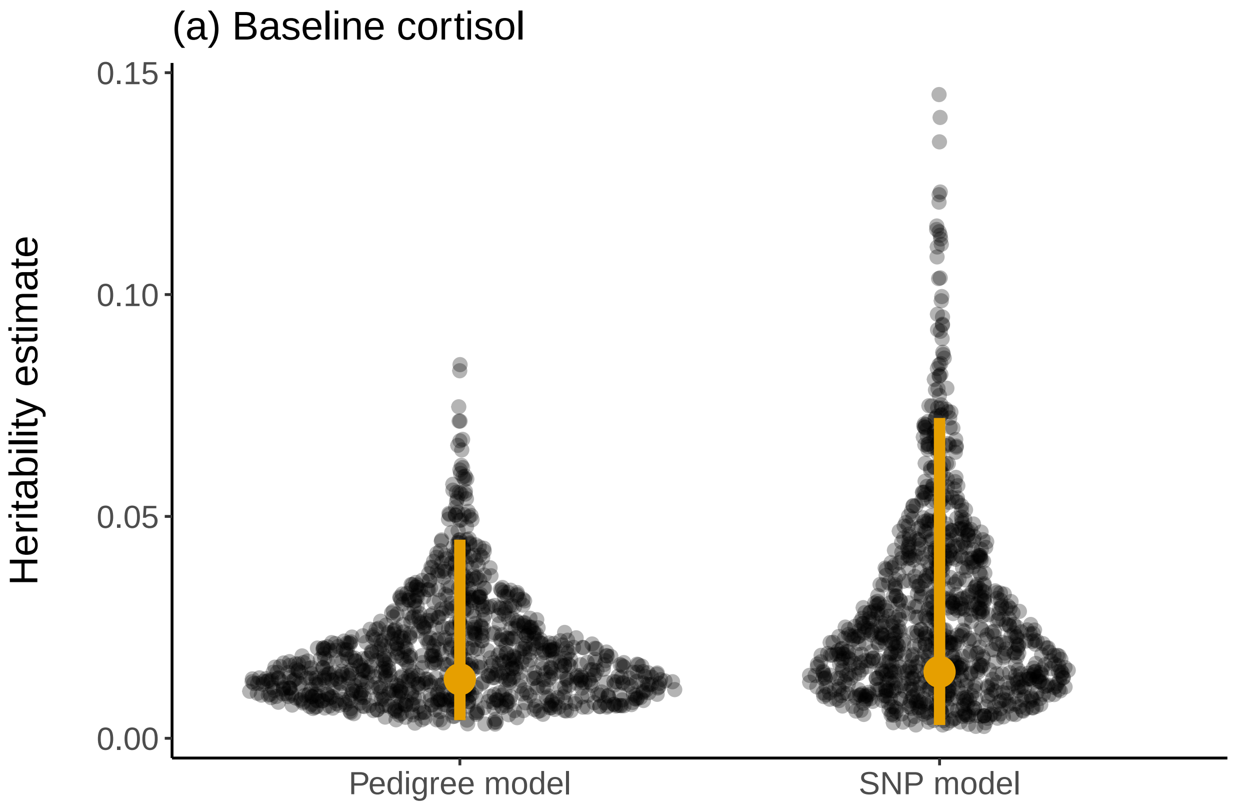

```
## - Session info ------------------------------------------------------------------
##  setting  value
##  version  R version 4.0.2 (2020-06-22)
##  os       Windows 10 x64
##  system   x86_64, mingw32
##  ui       RTerm
##  language (EN)
##  collate  English_United States.1252
##  ctype    English_United States.1252
##  tz       Europe/Berlin
##  date     2022-02-04
##
## - Packages ----------------------------------------------------------------------
##  package      * version    date       lib source
##  abind          1.4-5      2016-07-21 [1] CRAN (R 4.0.0)
##  ape          * 5.4-1      2020-08-13 [1] CRAN (R 4.0.2)
##  assertthat     0.2.1      2019-03-21 [1] CRAN (R 4.0.2)
##  backports      1.1.10     2020-09-15 [1] CRAN (R 4.0.2)
##  bayestestR     0.7.2      2020-07-20 [1] CRAN (R 4.0.2)
##  blob           1.2.1      2020-01-20 [1] CRAN (R 4.0.2)
##  boot           1.3-25     2020-04-26 [1] CRAN (R 4.0.3)
##  broom          0.7.0      2020-07-09 [1] CRAN (R 4.0.2)
##  car          * 3.0-9      2020-08-11 [1] CRAN (R 4.0.2)
##  carData      * 3.0-4      2020-05-22 [1] CRAN (R 4.0.0)
##  cellranger     1.1.0      2016-07-27 [1] CRAN (R 4.0.2)
##  cli            2.0.2      2020-02-28 [1] CRAN (R 4.0.2)
##  coda         * 0.19-3     2019-07-05 [1] CRAN (R 4.0.2)
##  codetools      0.2-16     2018-12-24 [2] CRAN (R 4.0.2)
##  colorspace     1.4-1      2019-03-18 [1] CRAN (R 4.0.2)
##  combinat       0.0-8      2012-10-29 [1] CRAN (R 4.0.3)
##  corpcor        1.6.9      2017-04-01 [1] CRAN (R 4.0.0)
##  crayon         1.3.4      2017-09-16 [1] CRAN (R 4.0.2)
##  cubature       2.0.4.1    2020-07-06 [1] CRAN (R 4.0.2)
##  curl           4.3        2019-12-02 [1] CRAN (R 4.0.2)
##  data.table   * 1.13.0     2020-07-24 [1] CRAN (R 4.0.2)
##  DBI            1.1.0      2019-12-15 [1] CRAN (R 4.0.2)
##  dbplyr         1.4.4      2020-05-27 [1] CRAN (R 4.0.2)
##  DHARMa       * 0.3.3.0    2020-09-08 [1] CRAN (R 4.0.2)
##  digest         0.6.25     2020-02-23 [1] CRAN (R 4.0.2)
##  dplyr        * 1.0.2      2020-08-18 [1] CRAN (R 4.0.2)
##  ellipsis       0.3.1      2020-05-15 [1] CRAN (R 4.0.2)
##  emmeans      * 1.5.0      2020-08-18 [1] CRAN (R 4.0.2)
##  estimability   1.3        2018-02-11 [1] CRAN (R 4.0.0)
##  evaluate       0.14       2019-05-28 [1] CRAN (R 4.0.2)
##  fansi          0.4.1      2020-01-08 [1] CRAN (R 4.0.2)
##  farver         2.0.3      2020-01-16 [1] CRAN (R 4.0.2)
##  fitdistrplus * 1.1-3      2020-12-05 [1] CRAN (R 4.0.3)
##  forcats      * 0.5.0      2020-03-01 [1] CRAN (R 4.0.2)
##  foreach        1.5.0      2020-03-30 [1] CRAN (R 4.0.2)
##  foreign        0.8-80     2020-05-24 [2] CRAN (R 4.0.2)
##  formatR      * 1.7        2019-06-11 [1] CRAN (R 4.0.2)
##  fs             1.5.0      2020-07-31 [1] CRAN (R 4.0.2)
##  gdata        * 2.18.0     2017-06-06 [1] CRAN (R 4.0.2)
##  generics       0.0.2      2018-11-29 [1] CRAN (R 4.0.2)
```

```
##  genetics       1.3.8.1.3  2021-03-01 [1] CRAN (R 4.0.4)
##  GeneticsPed    1.50.0     2020-04-27 [1] Bioconductor
##  ggforce      * 0.3.2      2020-06-23 [1] CRAN (R 4.0.3)
##  ggplot2      * 3.3.2      2020-06-19 [1] CRAN (R 4.0.2)
##  ggpubr       * 0.4.0      2020-06-27 [1] CRAN (R 4.0.2)
##  ggsignif       0.6.0      2019-08-08 [1] CRAN (R 4.0.2)
##  glue           1.4.2      2020-08-27 [1] CRAN (R 4.0.2)
##  goft         * 1.3.6      2020-06-25 [1] CRAN (R 4.0.3)
##  gtable         0.3.0      2019-03-25 [1] CRAN (R 4.0.2)
##  gtools         3.8.2      2020-03-31 [1] CRAN (R 4.0.0)
##  haven          2.3.1      2020-06-01 [1] CRAN (R 4.0.2)
##  hms            0.5.3      2020-01-08 [1] CRAN (R 4.0.2)
##  htmltools      0.5.0      2020-06-16 [1] CRAN (R 4.0.2)
##  httr           1.4.2      2020-07-20 [1] CRAN (R 4.0.2)
##  igraph         1.2.5      2020-03-19 [1] CRAN (R 4.0.2)
##  insight        0.9.5      2020-09-07 [1] CRAN (R 4.0.2)
##  iterators      1.0.12     2019-07-26 [1] CRAN (R 4.0.2)
##  jsonlite       1.7.1      2020-09-07 [1] CRAN (R 4.0.2)
##  kableExtra   * 1.2.1      2020-08-27 [1] CRAN (R 4.0.2)
##  knitr        * 1.29       2020-06-23 [1] CRAN (R 4.0.2)
##  labeling       0.3        2014-08-23 [1] CRAN (R 4.0.0)
##  lattice        0.20-41    2020-04-02 [2] CRAN (R 4.0.2)
##  lifecycle      0.2.0      2020-03-06 [1] CRAN (R 4.0.2)
##  lme4         * 1.1-27.1   2021-06-22 [1] CRAN (R 4.0.5)
##  lubridate      1.7.9      2020-06-08 [1] CRAN (R 4.0.2)
##  magrittr       1.5        2014-11-22 [1] CRAN (R 4.0.2)
##  MASS         * 7.3-52     2020-08-18 [2] CRAN (R 4.0.2)
##  Matrix       * 1.2-18     2019-11-27 [2] CRAN (R 4.0.2)
##  MCMCglmm     * 2.32       2021-03-12 [1] CRAN (R 4.0.4)
##  mgcv         * 1.8-33     2020-08-27 [2] CRAN (R 4.0.2)
##  minqa          1.2.4      2014-10-09 [1] CRAN (R 4.0.2)
##  mnormt         2.0.2      2020-09-01 [1] CRAN (R 4.0.2)
##  modelr         0.1.8      2020-05-19 [1] CRAN (R 4.0.2)
##  multcomp       1.4-13     2020-04-08 [1] CRAN (R 4.0.2)
##  MuMIn        * 1.43.17    2020-04-15 [1] CRAN (R 4.0.2)
##  munsell        0.5.0      2018-06-12 [1] CRAN (R 4.0.2)
##  mvtnorm        1.1-1      2020-06-09 [1] CRAN (R 4.0.0)
##  nlme         * 3.1-149    2020-08-23 [2] CRAN (R 4.0.2)
##  nloptr         1.2.2.2    2020-07-02 [1] CRAN (R 4.0.2)
##  numDeriv       2016.8-1.1 2019-06-06 [1] CRAN (R 4.0.0)
##  openxlsx       4.1.5      2020-05-06 [1] CRAN (R 4.0.2)
##  patchwork    * 1.1.0      2020-11-09 [1] CRAN (R 4.0.3)
##  performance  * 0.5.0      2020-09-12 [1] CRAN (R 4.0.2)
##  pillar         1.4.6      2020-07-10 [1] CRAN (R 4.0.2)
##  pkgconfig      2.0.3      2019-09-22 [1] CRAN (R 4.0.2)
##  plyr         * 1.8.6      2020-03-03 [1] CRAN (R 4.0.2)
##  polyclip       1.10-0     2019-03-14 [1] CRAN (R 4.0.3)
##  purrr        * 0.3.4      2020-04-17 [1] CRAN (R 4.0.2)
##  R6             2.4.1      2019-11-12 [1] CRAN (R 4.0.2)
##  Rcpp           1.0.7      2021-07-07 [1] CRAN (R 4.0.5)
##  readr        * 1.3.1      2018-12-21 [1] CRAN (R 4.0.2)
##  readxl         1.3.1      2019-03-13 [1] CRAN (R 4.0.2)
##  reprex         0.3.0      2019-05-16 [1] CRAN (R 4.0.2)
##  reshape2     * 1.4.4      2020-04-09 [1] CRAN (R 4.0.2)
```

```
##  rio            0.5.16     2018-11-26 [1] CRAN (R 4.0.2)
##  rlang          0.4.8      2020-10-08 [1] CRAN (R 4.0.3)
##  rmarkdown      2.3        2020-06-18 [1] CRAN (R 4.0.2)
##  rstatix        0.6.0      2020-06-18 [1] CRAN (R 4.0.2)
##  rstudioapi     0.11       2020-02-07 [1] CRAN (R 4.0.2)
##  rvest          0.3.6      2020-07-25 [1] CRAN (R 4.0.2)
##  sandwich       2.5-1      2019-04-06 [1] CRAN (R 4.0.2)
##  scales         1.1.1      2020-05-11 [1] CRAN (R 4.0.2)
##  sessioninfo    1.1.1      2018-11-05 [1] CRAN (R 4.0.2)
##  sn           * 1.6-2      2020-05-26 [1] CRAN (R 4.0.3)
##  specr        * 0.2.1      2020-03-26 [1] CRAN (R 4.0.3)
##  stringi        1.5.3      2020-09-09 [1] CRAN (R 4.0.2)
##  stringr      * 1.4.0      2019-02-10 [1] CRAN (R 4.0.2)
##  survival     * 3.2-3      2020-06-13 [2] CRAN (R 4.0.2)
##  tensorA        0.36.1     2018-07-29 [1] CRAN (R 4.0.0)
##  TH.data        1.0-10     2019-01-21 [1] CRAN (R 4.0.2)
##  tibble       * 3.0.3      2020-07-10 [1] CRAN (R 4.0.2)
##  tidyr        * 1.1.2      2020-08-27 [1] CRAN (R 4.0.2)
##  tidyselect     1.1.0      2020-05-11 [1] CRAN (R 4.0.2)
##  tidyverse    * 1.3.0      2019-11-21 [1] CRAN (R 4.0.2)
##  tmvnsim        1.0-2      2016-12-15 [1] CRAN (R 4.0.0)
##  tweenr         1.0.1      2018-12-14 [1] CRAN (R 4.0.3)
##  vctrs          0.3.4      2020-08-29 [1] CRAN (R 4.0.2)
##  viridisLite    0.3.0      2018-02-01 [1] CRAN (R 4.0.2)
##  webshot        0.5.2      2019-11-22 [1] CRAN (R 4.0.2)
##  withr          2.3.0      2020-09-22 [1] CRAN (R 4.0.2)
##  xfun           0.17       2020-09-09 [1] CRAN (R 4.0.2)
##  xml2           1.3.2      2020-04-23 [1] CRAN (R 4.0.2)
##  xtable         1.8-4      2019-04-21 [1] CRAN (R 4.0.2)
##  yaml           2.2.1      2020-02-01 [1] CRAN (R 4.0.2)
##  zip            2.1.1      2020-08-27 [1] CRAN (R 4.0.2)
##  zoo            1.8-8      2020-05-02 [1] CRAN (R 4.0.2)
##
## [1] C:/Users/localadmin/Documents/R/win-library/4.0
## [2] C:/Program Files/R/R-4.0.2/library
```
