## Supplementary Material for "Low heritability and high phenotypic plasticity of salivary cortisol in response to environmental heterogeneity in a wild pinniped"

**Supplementary Material:
Low heritability and high phenotypic plasticity of cortisol in
response to environmental heterogeneity in fur seals**

Rebecca Nagel, Sylvia Kaiser, Claire Stainfield, Camille Toscani, Cameron Fox-Clarke, Anneke J. Paijmans, Camila Costa Castro, David L. J. Vendrami, Jaume Forcada, Joseph I. Hoffman

*Validation of enzyme-linked immunosorbent assays (cortisol free in saliva DES6611, Demeditec Diagnostics GmbH, Kiel, Germany) on the Antarctic fur seal.*

Table S1: To assess linearity, we ran a serial dilution of three pooled samples in duplicate.

|  | measured ng/ml | expected ng/ml | linearity % |
| --- | --- | --- | --- |
| **sample 1** | | | |
| neat | 1.27 |  |  |
| 1:2 | 0.63 | 0.64 | 99.2 |
| 1:4 | 0.31 | 0.32 | 97.6 |
| 1:8 | 0.17 | 0.16 | 107.1 |
| 1:16 | 0.08 | 0.0079 | 100.8 |
| **sample 2** | | | |
| neat | 3.5 |  |  |
| 1:2 | 1.54 | 1.75 | 88.0 |
| 1:4 | 0.77 | 0.88 | 88.0 |
| 1:8 | 0.46 | 0.44 | 105.1 |
| 1:16 | 0.22 | 0.219 | 100.6 |
| **sample 3** | | | |
| neat | 7.08 |  |  |
| 1:2 | 3.41 | 3.54 | 96.3 |
| 1:4 | 1.49 | 1.77 | 84.2 |
| 1:8 | 0.73 | 0.89 | 82.5 |
| 1:16 | 0.35 | 0.44 | 79.1 |

Table S2: To assess the recovery rate, three pooled samples of different concentrations were spiked with different amounts of cortisol and measured in duplicate.

|  | measured ng/ml | expected ng/ml | recovery rate % |
| --- | --- | --- | --- |
| **sample 1** | | | |
| initial value | 0.33 |  |  |
| + 0.2 ng/ml | 0.63 | 0.53 | 118 |
| + 0.85 ng/ml | 1.34 | 1.18 | 114 |
| + 3.5 ng/ml | 4.36 | 3.83 | 114 |
| **sample 2** | | | |
| initial value | 0.82 |  |  |
| + 0.2 ng/ml | 1.00 | 1.02 | 98 |
| + 0.85 ng/ml | 1.77 | 1.67 | 106 |
| + 3.5 ng/ml | 5.31 | 4.32 | 123 |
| **sample 3** | | | |
| initial value | 1.67 |  |  |
| + 0.2 ng/ml | 1.93 | 1.87 | 103 |
| + 0.85 ng/ml | 2.63 | 2.52 | 104 |
| + 3.5 ng/ml | 6.59 | 5.17 | 127 |

Table S3: Full generalized linear mixed models of (a) pup and (b) maternal baseline cortisol. Here, we tested for among-individual variability by comparing model fit with and without ID as a random effect.

| (a) | Fixed effects | Random effects | AIC | BIC | χ^2^ | *p* |
| --- | --- | --- | --- | --- | --- | --- |
| Pup baseline cortisol | Age +  Body condition +  Weight +  Sex +  Season * Beach | - | 1556.8 | 1589.9 |  |  |
|  |  | (1\|ID) | 1543.0 | 1579.7 | 15.842 | < 0.001 |
|  |  | (Age\|ID) | 1488.9 | 1532.9 | 58.130 | < 0.001 |
| (b) |  | | | | | |
| Maternal baseline cortisol | Age +  Body condition +  Weight +  Season * Beach | - | 732.68 | 747.49 |  |  |
|  |  | (1\|ID) | 689.08 | 715.87 | 36.604 | < 0.001 |

*Chloroform-isoamylalcohol protocol for DNA extraction*

We extracted total genomic DNA from tissue samples using a standard chloroform-isoamylalcohol protocol. Approximately 2 mm2 of tissue was digested overnight at 55°C in 500µl of extraction buffer (10mM Tris, 2mM EDTA, 10mM NaCl, 1% SDS) and 20µl of proteinase K (10mg/ml). The following day, total genomic DNA was extracted by adding 250µl 5M NaCl and 700µl cold choloroform-isoamylalcohol (Roti®-C/T). The mixture was centrifuged for ten minutes to separate the DNA from the proteins, RNA, and lipids. 500µl of the aqueous upper phase was transferred to a new tube, where 50µl of 3M NaAc and 300µl of isopropanol was added. The mixture was again centrifuged for 20 minutes, after which the supernatent was discarded. The DNA pellet was incubated at room temperature in 500µl 70% ethanol for ten minutes, before being centrifuged for five minutes. The ethanol was then discarded and the pellet was left to dry at room temperature for a minimum of 30 minutes. The DNA pellet was rehydrated in 50µl of elution buffer (1 x TE) and stored at -20°C until further processing.
